# ILK binding to β1 Integrin is indirectly mediated by Kindlin-2

**DOI:** 10.64898/2026.05.06.723202

**Authors:** Susanne C.M. Reinhardt, Ralph T. Böttcher, Florian Brod, Jan D. Speidel, Ralf Jungmann, Reinhard Fässler

## Abstract

Integrin-linked kinase (ILK) and kindlin-2 (K2) are key components of focal adhesions (FAs) that regulate cell-matrix adhesion and integrin signaling. Both proteins directly bind each other, but how they influence each other’s localization to FAs and binding to integrins remains a subject of ongoing debate. Here, we establish a sensitive workflow to study protein-protein interactions in cells by combining methods from biochemistry, cell biology and super-resolution microscopy. Together with an analytical framework this approach allowed us to distinguish direct from indirect molecular interactions and construct detailed interaction networks. Disrupting the ILK-K2 interaction reduced ILK localization to FAs and compromised integrin function, whereas K2 recruitment was unaffected. Our interdisciplinary approach also revealed that ILK does not directly bind β1-integrin cytosolic domains in vitro and in cells. Instead, ILK was recruited to integrins exclusively through a K2-dependent mechanism, primarily via K2 bridging ILK and β1 integrins. These findings define the hierarchical relationship between ILK and K2 in FAs and highlight the essential role of K2-mediated ILK recruitment for integrin adhesion and signaling.

**Significance Statement:** How cells anchor to their environment is a fundamental question in biology. Integrins provide such a connection by bridging the extracellular matrix and the cytoskeleton. A central regulator of the integrin machine is Integrin-linked kinase (ILK). How ILK is recruited to β1 integrins is hotly debated since its discovery more than 30 years ago. By integrating cell biology and biochemistry with super-resolution DNA-PAINT microscopy and a novel spatial analysis framework, we demonstrate that ILK does not bind directly to integrin cytoplasmic tails. Instead, we found that ILK is recruited by kindlin-2 (K2) to active, adhesion plaque-resident integrins. This work resolves a long-standing controversy in cell biology and establishes a versatile workflow for distinguishing direct from indirect protein-protein interactions *in situ*.

## Introduction

Integrins mediate adhesion between cells and to the extracellular matrix (ECM). A hallmark of integrins is that ligand binding requires an activation step, characterized by conformational changes affecting the entire molecule. This process is facilitated by talin and kindlin binding to the β-integrin cytoplasmic tail as well as transmission of forces from actomyosin to the integrin•ligand bonds(1). Upon activation and ligand binding, integrins recruit hundreds of ancillary proteins to the adhesion site, which is collectively termed the adhesome(2, 3) and bidirectionally transmits mechanical and biochemical signals to control cell survival, proliferation, migration, and differentiation. A key FA protein is integrin-linked kinase(4) (ILK), which forms a ternary complex with particularly interesting new cysteine-histidine-rich protein (PINCH) and parvin (IPP complex) and is believed to bind to β-integrin tails either directly or indirectly via kindlins (**Fig. 1a**).

**Figure 1.**
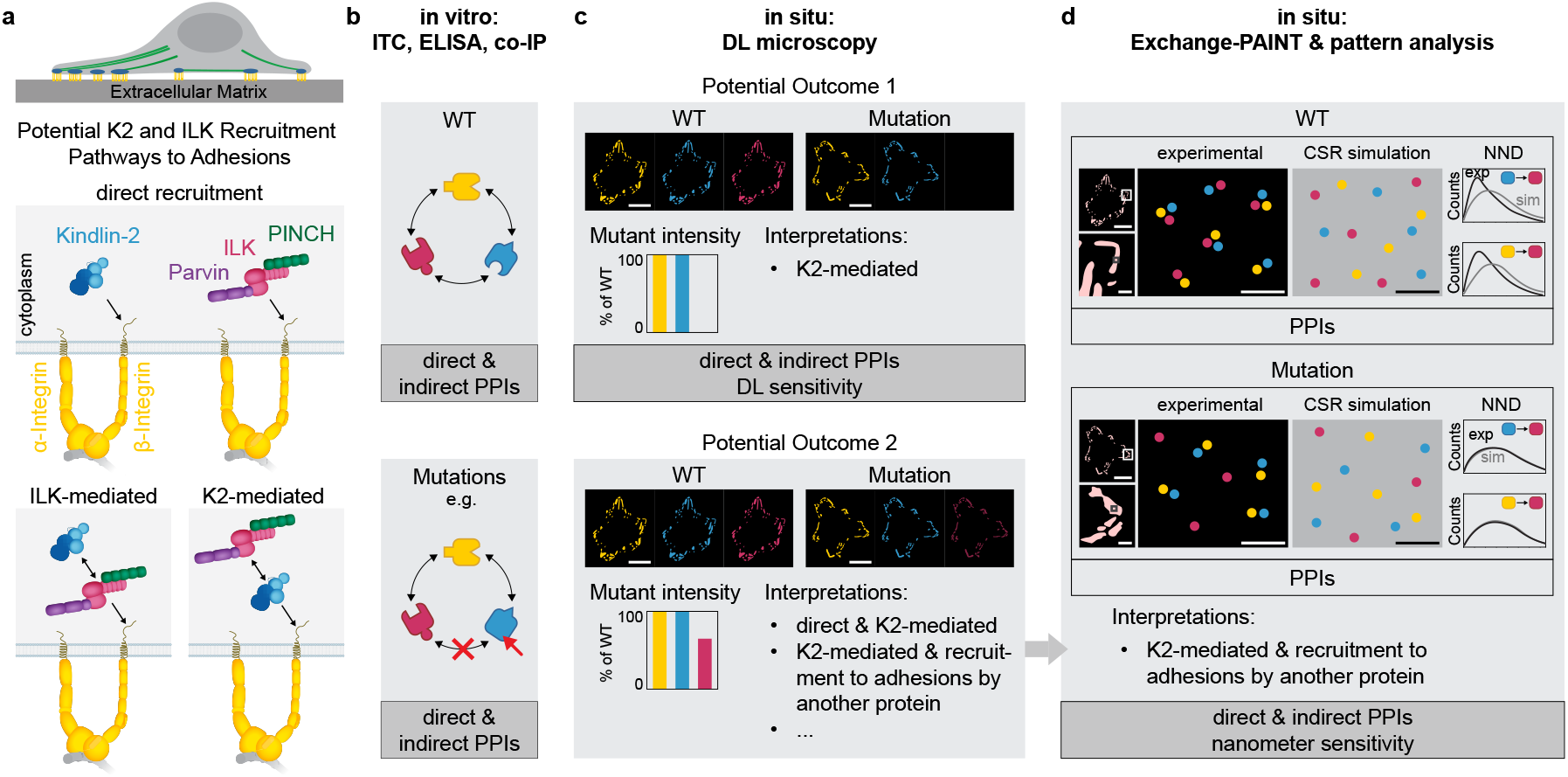
Assessing in situ protein-protein interaction patterns in FAs using diffraction-limited (DL) and Exchange-PAINT microscopy. **a** Integrin-mediated adhesions connect the actin cytoskeleton to the ECM. The integrin-linked kinase (ILK), PINCH, and parvin (IPP) complex, along with kindlin-2 (K2), are critical regulators of integrin function and signalling. Three potential mechanisms for K2 and ILK recruitment to FAs are proposed: (1) direct, independent binding to β-integrin cytoplasmic tails, (2) ILK-mediated recruitment of K2, or (3) K2-mediated recruitment of ILK. **b-d** Protein-protein interactions (PPIs) are studied in a three-step workflow. **b** First, the *in vitro* binding capability of WT proteins is confirmed, and binding-deficient mutations are evaluated. **c** Second, DL microscopy of WT and mutant cells is performed. If the comparison of protein localization and intensities permits only one single interpretation, *in situ* interaction patterns can be identified (e.g. potential outcome 1 representing complete loss of colocalization in a DL volume such as FAs). However, when multiple interpretations are possible, in situ PPIs cannot be inferred (e.g. potential outcome 2 showing a reduction of intensities). **d** Third, for ambiguous cases, multiplexed super-resolution microscopy with single molecule resolution e.g. Exchange-PAINT is combined with a pattern analysis framework and reference simulations to define recruitment pathways. This approach enables differentiation between direct and indirect interactions in situ at nanoscale resolution. Scale bars 20 µm (full cells), 5 µm (zoom-in to adhesions) and 50 nm (zoom-in to single molecule resolution).

The kindlin family consists of three FERM (protein 4.1, ezrin, radixin, moesin) domain-containing proteins (kindlin-1, -2, -3) with distinct expression patterns. Kindlins is among the earliest proteins recruited to the integrin β-tail, where it cooperates with talin to stabilize integrin-ligand bonds, induces assembly of nascent adhesions, and recruits additional proteins such as ILK, paxillin, etc. to initiate signalling. ILK was discovered through a yeast two-hybrid screen using the cytoplasmic tail of β1 integrin as bait(5), and was shown to co-immunoprecipitate with β1- and β3-integrin subunits using a polyclonal anti-ILK antiserum(6). However, several follow-up studies were unable to confirm a direct interaction of ILK with integrin β-tails. For instance, yeast two-hybrid studies with the highly conserved *Drosophila* ILK protein failed to detect an interaction with the highly conserved cytoplasmic domain of integrin β PS(7). Furthermore, mass spectrometry (MS)-based interactome experiments were also unable to demonstrate an interaction between mammalian ILK and β-integrin tails(8-11). Therefore, it was speculated that ILK is recruited to FAs not via binding to β1-integrin tails, but rather indirectly through FA protein(s).

ILK directly binds to kindlin, which was first reported in *C. elegans* and subsequently confirmed in mammals(12-18). The interaction between K2 and ILK occurs between the F2 subdomain of K2 and the pseudokinase domain (pKD) of ILK(16-19). The disruption of the ILK-K2 interaction in cells revealed conflicting results (**Fig. 1a**). Whereas one study reported reduced ILK recruitment to FAs, another described a more complex, co-dependent relationship between kindlin and ILK(15, 18). The latter is based on observations both in mammalian cells and *C. elegans* muscle cells *in vivo*, showing that the disruption of the ILK-kindlin interaction curbs kindlin recruitment to integrin adhesion sites(13, 16, 17, 19). How kindlin-binding-deficient ILK decreases kindlin levels in FAs has not been further studied.

To address these questions, we investigate how ILK recruitment to FAs relates to interactions between individual molecules within FAs. Establishing this connection requires an approach that measures *in situ* protein-protein interactions (PPIs) while providing spatial context with sensitivity ranging from molecular to subcellular length scales. Mass spectrometry (MS) of affinity purifications excels at mapping interaction networks and identifying novel binding partners. While MS(3) and cross-linking MS(2, 20) on purified FAs as well as proximity biotinylation (BioID) MS(21, 22) advanced our understanding of the adhesome, these techniques have limitations. They lack sensitivity to changes in protein abundance and cannot provide insight into protein proximity across the range of spatial scales relevant to adhesome recruitment. Protein ligation assays(23) (PLA) and Förster resonance energy transfer(24, 25) (FRET) detect proximity between molecules and reveal their diffraction-limited location in the cell, offering insights into molecular interactions. More recent approaches have achieved sub-diffraction localization of molecule pairs by employing split detection sites in super-resolution microscopy(26, 27). However, these methods are insensitive to changes in proximity beyond a method-dependent distance threshold and cannot reliably distinguish between proximity arising from specific interactions or random collisions.

Switching from a proximity-based fluorescent readout to mapping protein coordinates addresses both challenges. Among microscopy techniques that have demonstrated single-molecule resolution in dense structures(28-31), DNA points accumulation for imaging in nanoscale topography (DNA-PAINT)(32, 33) enables straightforward multiplexing through Exchange-PAINT(34) and provides sufficient throughput for whole-cell imaging with adequate statistics. These protein maps enable direct distance measurements at any length scale in the cell, with the lower boundary defined by the resolution of the employed technique. Importantly, they also provide essential data required to generate realistic simulations of protein distributions. By comparing observed protein colocalization patterns with reference simulations, colocalization mediated by interaction (of unknown nature) can be distinguished from random proximity. Peptide-(35) and DNA-PAINT(36) have provided maps of the FA protein talin and revealed its non-random colocalization with K2. However, while an indirect, integrin-mediated interaction appears likely, the exact nature of the interaction – direct or mediated by other molecules – remains inaccessible through imaging and simulation alone.

To answer this type of question, we integrate biochemically and cell-biologically verified WT and mutant PPI patterns into a workflow to identify direct and indirect PPI patterns in intact cells (**Fig. 1b, c**). By characterizing these interactions *in situ* using Exchange-PAINT microscopy at single-molecule resolution and employing a proximity-based analysis framework (**Fig. 1d**), we identified how ILK is recruited to FAs and integrins. More generally, this workflow offers a scalable and sensitive approach for PPI studies, adaptable across diverse cellular systems and complex interaction networks.

## Results

### Recombinant full-length IPP complex does not bind β1 and β3 Integrin cytosolic tails

Previous studies typically used truncated versions of IPP complex members to recombinantly express and purify a minimal heterotrimeric IPP complex(6, 37). For biochemical and biophysical experiments, we utilised a multigene baculovirus expression system and purification scheme to obtain an IPP complex containing full-length murine ILK, PINCH1 and β-parvin. This approach yielded a soluble, stable and monodisperse complex in solution (**Fig. S1a, b**). The size of ILK, β-parvin and PINCH1 on SDS-PAGE and by whole mass proteomics is consistent with their expected molecular weights of ∼51 kDa, ∼45 kDa, and ∼38 kDa, respectively (**Fig. S1a**). Furthermore, the purified IPP complex is resolved in a single peak by mass photometry corresponding to a molecular weight of approximately 129 kDa, in good agreement with the expected value of ∼134 kDa (**Fig. S1b**).

The IPP complex associates with several adhesion proteins that potentially facilitate its recruitment to FAs, most notably kindlin-1 and -2 (13, 16-18), paxillin (38-43) and cytoplasmic tails of β1 and β3 integrins (5, 6). We therefore asked whether these interactions can be recapitulated *in vitro* with our full-length recombinant IPP complex. Mass photometry measurements revealed that the addition of recombinant K2 led to the formation of IPP/K2 complexes represented by an additional peak with an average mass of around 207 kDa (**Fig. 2a**). Mass photometry was not used to analyse IPP complexes with integrin tail peptides, as the low molecular weight of the peptides prevents clear separation of the peaks corresponding to IPP alone and IPP bound to peptide. Therefore, we used isothermal titration calorimetry (ITC), which revealed no interaction between the full-length IPP complex with either β1 or β3 cytoplasmic tail peptides (**Fig. 2b**). To corroborate these findings, we performed an enzyme-linked immunosorbent assay (ELISA), which confirmed that neither β1 nor β3 cytoplasmic tail peptides bind to the IPP complex (**Fig. 2d-f** and **Fig. S1c**). The ELISA assay also revealed that wild-type K2 (K2-WT) readily bound to wild-type β1- and β3-tail peptides (β1-WT, β3-WT), but not to scrambled peptides (β1-Scr, β3-Scr) or β1-peptides with mutations in the kindlin binding site (β1-YY/AA) (**Fig. 2c, e** and **Fig. S1c**). To test whether IPP binding to integrin tails depends on K2, we incubated β1-WT-tail or control β1-tail peptides (β1-Scr or β1-YY/AA) with either the IPP complex alone or in combination with purified K2-WT, integrin-binding-deficient K2 (K2 Q614A W615A; K2-QW/AA), or ILK-binding-deficient K2 (K2 L353A L357A; K2-LL/AA)(16). K2-WT and K2-LL/AA showed comparable binding to β1-WT peptides, while K2-QW/AA failed to interact with β1-WT-tail peptides (**Fig. 2f** and **Fig. S1d**). Notably, both β1- and β3-WT-tail peptides readily bound the IPP complex in the presence of K2-WT, while the β1-WT-tail peptide was unable to recruit the IPP complex in the presence of the integrin- or ILK-binding deficient K2 (**Fig. 2f, Fig. S1e**). While IPP binding to integrin tail peptides was K2 dependent, K2 binding to β1-WT tail peptide occurred independent of the IPP complex (**Fig. S1f**). Together, these data indicate that the IPP complex binds to the β1 and β3 cytoplasmic tail through K2 *in vitro* and requires K2 binding to integrin and ILK.

**Figure 2.**
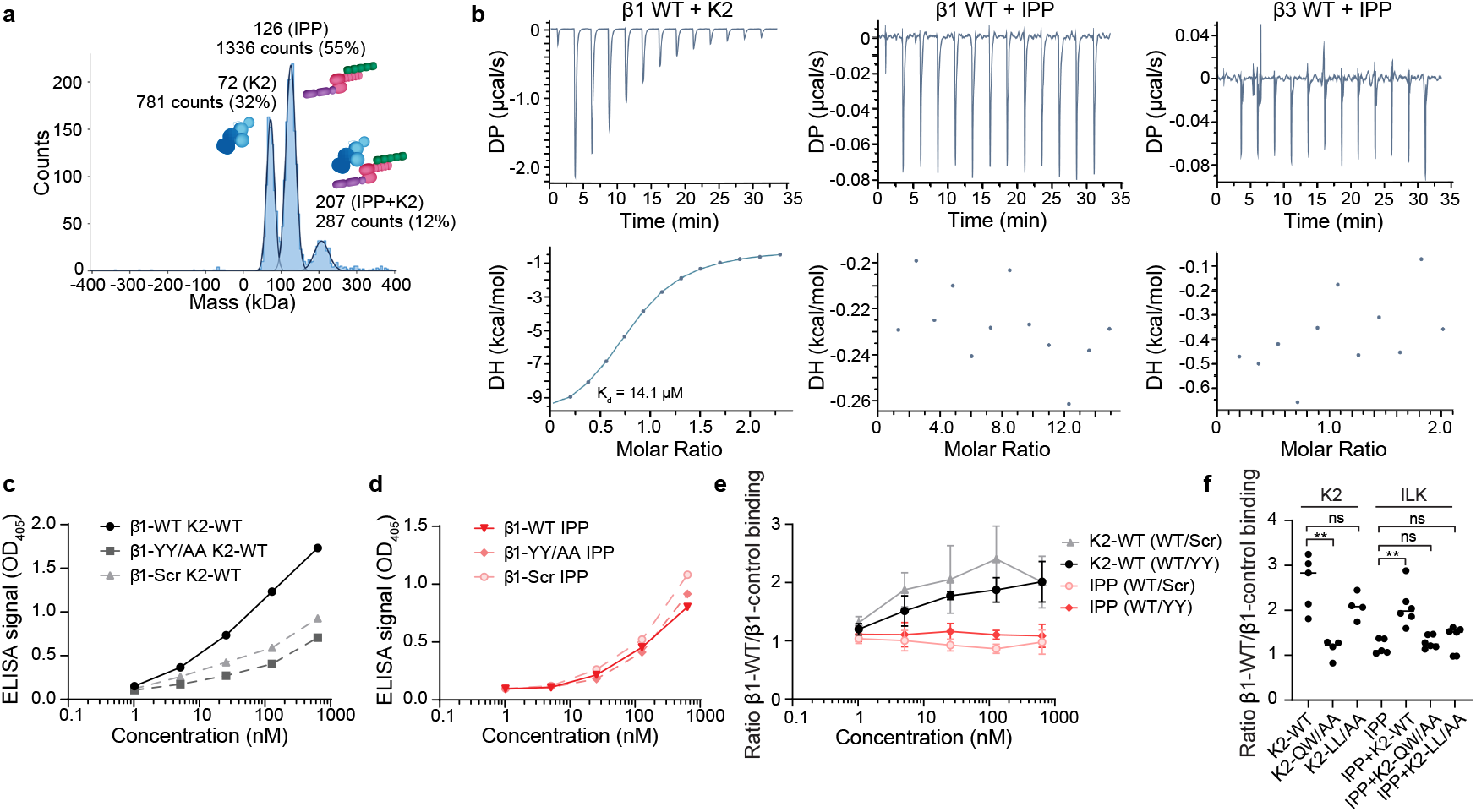
IPP binding to β1 and β3 integrin cytoplasmic tails is K2 dependent. **a** Analysis of the IPP-K2 interaction by mass photometry. Histogram of mass distributions observed for a 1:1 molar mixture of IPP complex and K2. **b** Isothermal titration calorimetry measurements of β1- and β3-tail peptides with K2 or IPP complex. Representative ITC curves of the interactions are shown. The top half of each panel shows the calorimetric titration of the integrin tail with the indicated ligand, and the lower half displays the integrated injection heats. K_d_ = 14.1 µM for β1-tail interaction with K2. **c**,**d** ELISA-based affinity measurements and quantification of wild-type K2 (c) and IPP complex (d) to the immobilized β1-WT, β1-YY/AA, or scrambled β1-tail peptides to determine nonspecific binding. Bound proteins were detected with antibodies against K2 (c) or ILK (d). Representative binding curves are shown. **e** Quantification of the relative binding of K2-WT or IPP to β1-WT tail peptides compared to β1-YY/AA or scrambled β1-tail peptides over the indicated concentration range (n=3 independent experiments; error bars indicate SD). **f** Increased binding of the IPP complex to β1-tail peptides after co-incubation with wild-type K2 but not integrin- or ILK-binding-deficient K2 (control β1-tail peptides: β1-YY/AA and scrambled β1). 25 nM IPP complex was applied to immobilized β1-WT, β1-YY/AA, or scrambled β1-tail peptides either alone or in combination with 250 nM K2-WT, K2-QW/AA or K2-LL/AA protein. After removing unbound proteins, remaining proteins were detected with antibodies against K2 or ILK (n=4-6 independent experiments, *t*-test significances are indicated: ns = not significant, ** 0.001 < p ≤ 0.01).

### IPP-K2 interaction efficiently recruits the IPP complex to FAs

To test if the interaction with K2 is required to recruit ILK to FAs in cells, we depleted ILK in talin-1/2- and kindlin-1/-2-deficient fibroblasts (qKO)(44) by CRISPR/Cas9 to obtain qKO cells lacking ILK expression (qKO/ILK-KO). Subsequently, two qKO/ILK-KO clones were reconstituted with mouse talin-1 and different combinations of N-terminally mCherry-tagged wild-type ILK (mCherry-ILK-WT) or kindlin-binding deficient mCherry-ILK K423D/I427E (mCherry-ILK-KI/DE)(18), and N-terminally mEGFP-tagged wild-type K2 (mEGFP-K2-WT) or ILK-binding deficient K2 (mEGFP-K2-L353A/L357A (mCherry-K2-LL/AA))(16) (**Fig. 3a** and **Fig. S2a**). Co-immunoprecipitation assays with these engineered cells revealed that mEGFP-K2-WT co-precipitated mCherry-ILK-WT, while mCherry-ILK-KI/DE or mEGFP-K2-LL/AA strongly reduced K2/ILK interaction (**Fig. 3b**). As expected, re-expression of mEGFP-K2-WT and mCherry-ILK-WT allowed the cells to adhere and spread on FN-coated surfaces. Interestingly, re-expression of the K2-binding-deficient mCherry-ILK-KI/DE or ILK-binding-deficient mEGFP-K2-LL/AA mutants only weakly impaired cell adhesion and spreading (**Fig. 3c, d** and **Fig. S2b**). We also quantified and characterized the FAs (**Fig. S3**) and found that the FA number per cell decreased and the FA area increased in mutant cells, while their aspect ratio and peripheral localization remained largely unaltered. To test whether the interaction with K2 is required for ILK recruitment to FAs, we visualized the subcellular localization of mCherry-ILK and mEGFP-K2 in cells by confocal microscopy. Expectedly, mEGFP-K2-WT localized to FAs. Interestingly, ILK-binding-deficient mEGFP-K2-LL/AA showed context-dependent localization. Whereas mEGFP-K2-LL/AA readily localized to FAs of kindlin-depleted cells (**Fig. 3e, f** and **Fig. S2c, d**), mEGFP-K2-LL/AA was markedly reduced at FAs of cells expressing endogenous K2 (**Fig. S2e**). This finding suggests that FA recruitment of K2 does not depend on its interaction with ILK and that wild-type K2 outcompetes ILK-binding-deficient K2 recruitment to FAs. In contrast, mCherry-tagged ILK and the IPP complex member α-parvin were only efficiently targeted to FAs when the ILK-binding-competent K2-WT was expressed (**Fig. 3e, f and Fig. S2c, d, f**). Collectively, these findings indicate that ILK relies on the interaction with K2 for efficient recruitment to FAs and mediate integrin functions, while the localization of K2 to FAs does not require ILK.

**Figure 3.**
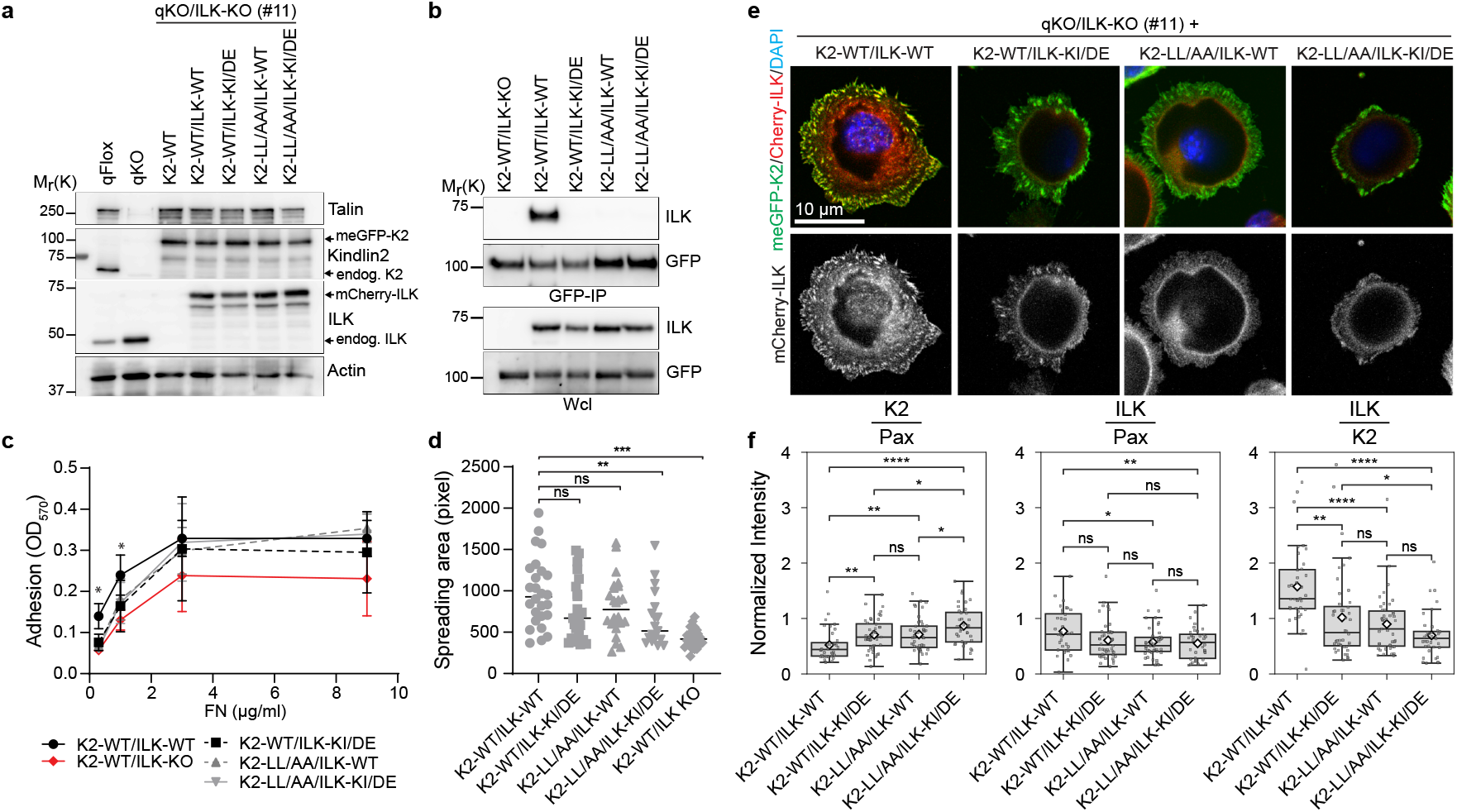
K2 is required for efficient ILK localization to FAs. **a** Western blot analysis of parental fibroblasts (qFlox), talin1/2 and kindlin1/2-deficient (qKO) cells, and ILK-deficient qKO cells (qKO/ILK-KO) reconstituted with talin-1 and different combinations of mCherry-tagged ILK (ILK-WT or ILK-KI/DE) and mEGFP-tagged K2 (K2-WT or K2-LL/AA). The bands for endogenous (endog.) ILK and K2 are indicated. Actin served as loading control. **b** Western blot of GFP-immunoprecipitations and whole cell lysates (wcl) from the indicated mEGFP-K2 and mCherry-ILK expressing cell lines. Blots were probed with antibodies against GFP and ILK. **c** Quantification of adhesion of indicated cell lines after seeding 30 min to increasing FN concentrations (n=4 independent experiments; error bars indicate SEM, *t*-test significances are calculated in correspondence to K2-WT/ILK-WT cells; a single asterisk (*) indicates that all comparisons at this concentration are significant, * 0.01 < p ≤ 0.05). **d** Quantification of the cell area of indicated cell lines after spreading on FN for 30 min (>50 cells counted in three independent experiments; *t*-test significances are indicated, ns = not significant, ** 0.001 < p ≤ 0.01, *** 0.0001 < p ≤ 0.001). **e** Fluorescence images of mEGFP-K2 and mCherry-ILK in kindlin and ILK-deficient (qKO/ILK-KO) cells allowed to spread for 30 min on FN. DAPI was used to stain nuclei. **f** FA-localized fluorescence intensities of mEGFP-K2 and mCherry-ILK were measured for cells seeded for 75 min on FN. Paxillin served as adhesion marker. K2 intensities were normalized with respect to paxillin. ILK intensities were normalized relative to paxillin or to K2. N ≥ 64 cells per condition were analysed. Welch’s *t*-test was applied to normalized intensities: ns (not significant) p > 0.05, * 0.01 < p ≤ 0.05, ** 0.001 < p ≤ 0.01, **** p ≤ 0.0001.

### Paxillin-independent pathways dominate IPP recruitment to FAs

ILK recruitment to FAs was reduced but not abolished in cells expressing the K2-LL/AA mutant, suggesting that the K2-LL/AA mutation only partially impairs the interaction, with ILK, or that alternative, K2-independent recruitment pathways enable ILK localization to FAs. FAs are complex and dynamic protein assemblies governed by intricate and multifaceted interactions. Paxillin emerges as a key bridging protein for an alternative pathway for ILK recruitment, as it directly interacts with both ILK and K2(39, 44-46) positioning it as a mediator that may compensate for disrupted direct K2-ILK interactions (**Fig. 4a**).

**Figure 4.**
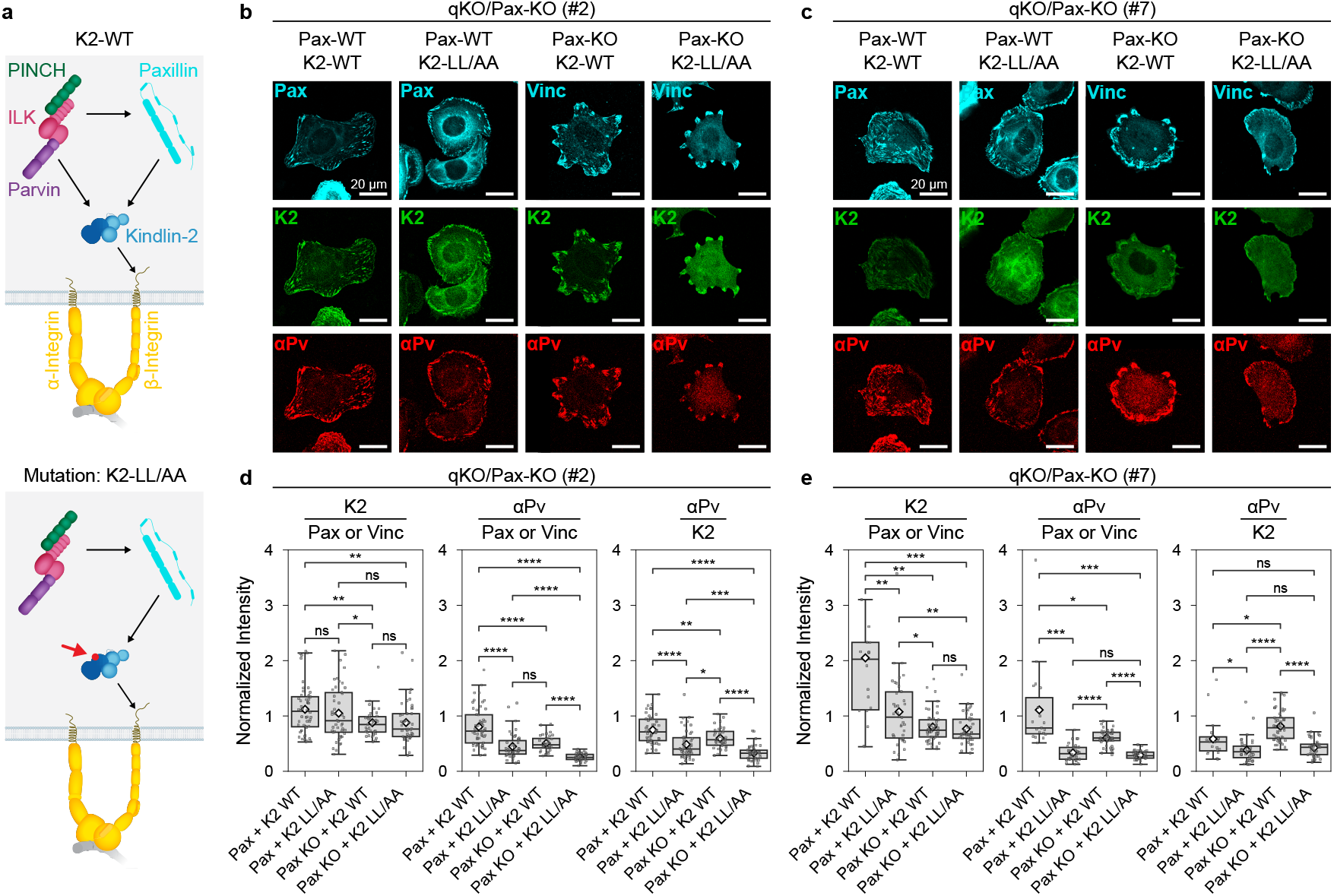
Paxillin plays a minor role in IPP recruitment to FAs. **a** Proposed paxillin-dependent recruitment of the IPP complex to integrin tails in the absence of a direct K2/ILK interaction. **b, c** Fluorescence images show mEGFP-tagged-K2-WT or mEGFP-tagged-K2-LL/AA and mCherry-paxillin in kindlin and paxillin-deficient (qKO/Pax-KO) cells for two independent clones. Cells were allowed to spread for 75 min on FN and stained with antibodies against endogenous α-parvin and vinculin. **d, e** Quantification of K2 and α-parvin recruitment to FAs as FA-localized fluorescence intensities of K2 and α-parvin. K2 intensities were normalized with respect to the adhesion markers paxillin and vinculin. α-parvin intensities were normalized relative to the adhesion markers or to K2. Notably, paxillin depletion did not consistently affect α-parvin intensities in K2-WT expressing cells but reduced α-parvin levels in FAs of K2-LL/AA expressing cells of clone 2. N ≥ 36 cells per condition were analysed. Welch’s *t*-test was applied to normalized α-parvin intensities: ns (not significant) p > 0.05, * 0.01 < p ≤ 0.05, ** 0.001 < p ≤ 0.01, *** 0.0001 < p ≤ 0.001, **** p ≤ 0.0001. Contrast settings were adjusted for each protein but kept constant across all cell lines.

To systematically investigate paxillin’s role in ILK recruitment, we depleted paxillin in talin1/2- and kindlin-1/2-deficient fibroblasts (qKO) using CRISPR/Cas9 technology. Two independent qKO/Pax-KO cell clones were subsequently reconstituted with mouse talin-1 and either mEGFP-K2-WT or mEGFP-K2-LL/AA, and either with or without mCherry-tagged paxillin. Confocal microscopy demonstrated that the mEGFP-K2-LL/AA mutation permitted K2 localization to FAs independent of the re-expression of paxillin (**Fig. 4b, c**). Interestingly, the quantification of fluorescent intensities in FAs of paxillin-deficient cells (**Fig. 4d, e**) revealed some inter-clonal variability: while clone 2 showed only a minor reduction of K2 intensity, clone 7 exhibited a pronounced decrease of K2 intensity in FAs. To account for these variations in K2 levels, we quantified α-parvin, an endogenous member of the IPP complex, relative to paxillin, vinculin and K2. Consistent with our previous observations (**Fig. 3e, f**), α-parvin recruitment was significantly impaired by the mEGFP-K2-LL/AA mutation in paxillin-deficient but also in paxillin re-expressing cells. In presence of K2-WT, paxillin depletion had no discernible effect on α-parvin recruitment. Notably, in K2-LL/AA reconstituted qKO/Pax-KO cells, the absence of paxillin led to a further reduction in α-parvin levels at FAs in clone 2, whereas clone 7 showed no additional decrease. Given that α-parvin remained detectable in FAs of mEGFP-K2-LL/AA expressing cells even in the absence of paxillin, we overall conclude that IPP complex recruitment to FAs is significantly more dependent on K2 than on paxillin, which is not essential and at most plays a minor role.

### Exchange-PAINT shows disruption of ILK interaction to K2-LL/AA *in situ*

Our confocal microscopy measurements could not unambiguously identify *in situ* recruitment pathways of ILK to FAs. To distinguish random ILK distribution in FAs from recruitment into the vicinity of K2, we enhanced the detection sensitivity and spatial resolution by employing super-resolution microscopy. We first characterized the association of ILK with K2-WT and K2-LL/AA using multiplexed Exchange-PAINT (**Fig. 5a**). To this end, mCherry-ILK and mEGFP-K2 were labelled with orthogonal DNA barcodes (“docking strands”) via DNA-conjugated anti-mCherry and anti-GFP nanobodies. Visualization of ILK and K2 was then achieved through sequential rounds of imaging with complementary dye-labelled “imager” strands. Their transient hybridization to the docking strands produces the blinking necessary for single-molecule localization microscopy.

**Figure 5.**
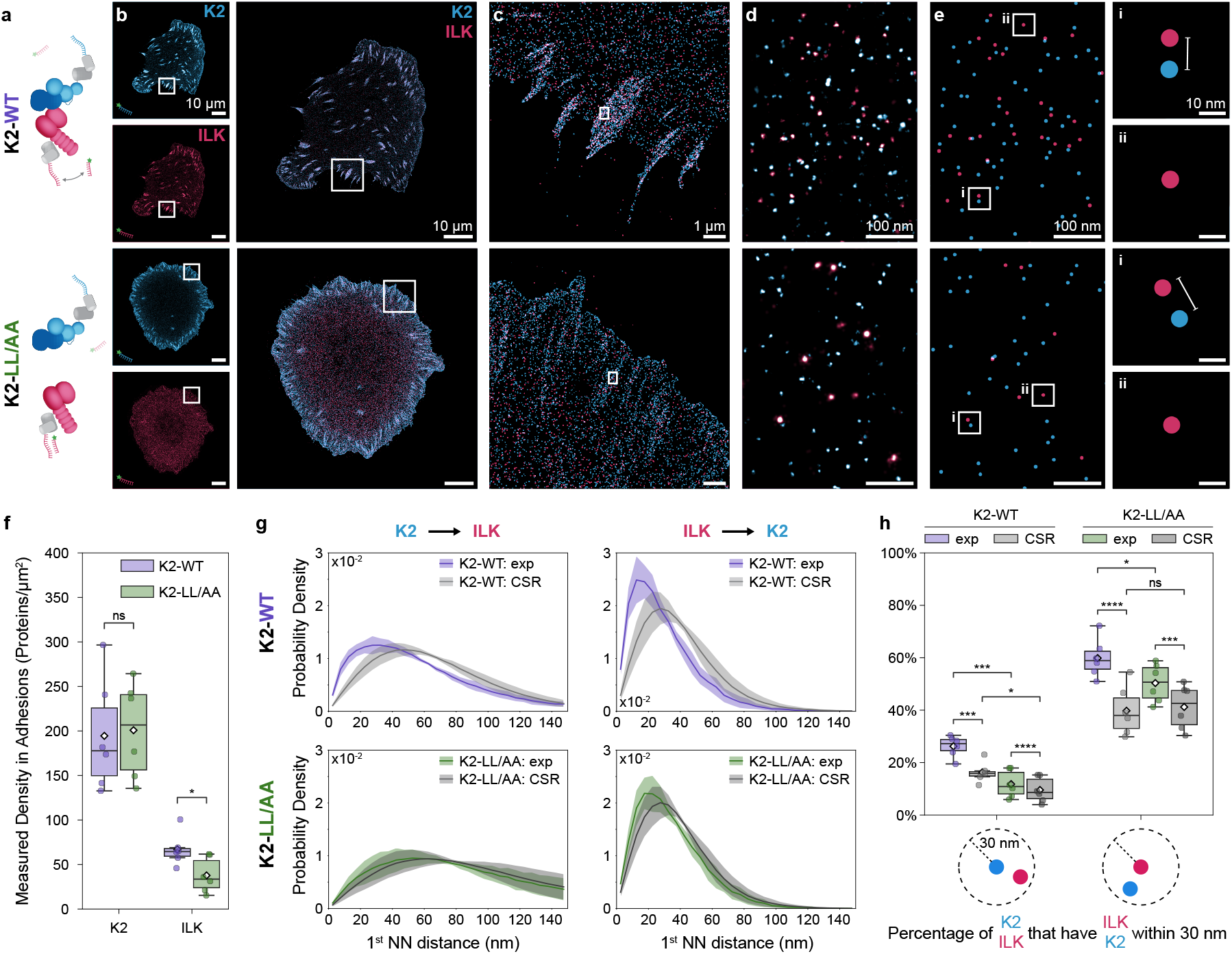
Exchange-PAINT demonstrates direct *in situ* binding between K2-WT and ILK, which is disrupted by the K2-LL/AA mutation. **a** Target proteins mEGFP-K2 and mCherry-ILK are labelled with orthogonal DNA docking strands via anti-mCherry and anti-GFP nanobodies. **b** Two-plexed Exchange-PAINT of representative cells expressing mCherry-ILK together with either mEGFP-K2-WT (top) or mEGFP-K2-LL/AA (bottom). **c** Magnified view of FAs. **d** Zoom into FAs. Spots correspond to individual mCherry-ILK and mEGFP-K2 molecules. **e** Protein coordinates were calculated and rendered as dots. The protein map reveals ILK - K2 (WT or LL/AA) pairs (e.g. regions i) as well as isolated ILK molecules (e.g. regions ii). **f** Observed surface density of K2 and ILK molecules in FAs of N = 6 cells per cell line. The density of K2 remains unaffected in K2-LL/AA expressing cells, while ILK density is significantly reduced in K2-LL/AA. Reported values represent observed densities and do not account for labelling efficiency. Welch’s t-test at α = 0.05; ns (not significant) p > 0.05, * 0.01 < p ≤ 0.05. **g** Average NND histograms between K2 and ILK in FAs compared to simulations of complete spatial randomness (CSR) (Mean ± SD of N = 6 cells per cell line). Left (K2 → ILK): The experimental distribution differs from the CSR scenario in K2-WT cells (p < 0.0001), but not in K2-LL/AA cells (p = 0.486). The experimental distribution in K2-LL/AA mutant cells is significantly shifted towards larger distances compared to WT (p < 0.0001). CSR distributions differ between cell lines (p < 0.0001). Right (ILK → K2): Experimental distributions differ significantly from CSR in K2-WT (p < 0.0001) and K2-LL/AA cells (p < 0.0001), while the K2-LL/AA is more similar to CSR. Experimental distributions between cell lines are shifted toward larger distances in the K2-LL/AA cells (p < 0.0001). The CSR distributions remain undistinguishable (p = 1). Modified Chi-squared test(56) at α = 0.05. **h** Percentage of K2 molecule with at least one ILK molecule within 30 nm (left) and vice versa (right) in FAs compared to CSR. The threshold of 30 nm accommodates the linkage error of nanobody labelling for mCherry-ILK and mEGFP-K2 (∼ 5-10 nm), alongside an average localization precision of ∼ 3 nm. In K2-LL/AA cells, both conditions are significantly reduced compared to WT. While both K2-LL/AA experimental distributions shift toward their respective CSR levels, they remain significantly above random expectation. Notably, the percentage of ILK molecules with K2 within 30 nm in the K2-LL/AA cell line remains further from its CSR baseline compared to the K2 with ILK case. Boxplots show median and 25th and 75th percentile with whiskers reaching the last data point within 1.5 × interquartile range, diamonds indicate mean. Paired two-sample t-test for exp vs. CSR and Welch’s t-test for K2-WT vs. K2-LL/AA. ns (not significant) p > 0.05, * 0.01 < p ≤ 0.05, *** 0.0001 < p ≤ 0.001, **** p ≤ 0.0001.

Consistent with our confocal microscopy results, Exchange-PAINT revealed FAs enriched in mEGFP-K2-WT and mEGFP-K2-LL/AA, whereas FA recruitment of mCherry-ILK-WT was visibly impaired in cells co-expressing mEGFP-K2-LL/AA compared to those co-expressing mEGFP-K2-WT (**Fig. 5b, c**). To quantify these observations, we identified the positions of individual proteins within each cell as centres of DNA-PAINT localization clusters (**Fig. 5d, e**, Methods) and subsequently defined FA regions using a binary mask (Methods and **Fig. S4**). This analysis corroborated our visual findings, recovering equivalent densities of K2-WT and K2-LL/AA in FAs, but a significant reduction of ILK densities in K2-LL/AA cells compared to K2-WT cells (**Fig. 5f** and **Fig. S5**). The protein maps revealed adjacent K2 and ILK molecules (**Fig. 5e** region **i**) as well as ILK proteins without an immediate K2 neighbour (**Fig. 5e** region **ii**) in both K2-WT and K2-LL/AA FAs. Notably, pairs of K2-LL/AA and ILK are observable despite the mutation, but appear less frequent than pairs of K2-WT and ILK.

To quantitatively assess the impact of K2-LL/AA on molecular proximity to ILK in FAs, we performed nearest neighbour distance (NND) analysis. The average NNDs in both directions (K2 → ILK and ILK → K2) showed greater separation between K2-LL/AA and ILK compared to K2-WT and ILK in FAs (**Fig. 5g**). To disentangle the effects of protein density from specific molecular interactions, we compared the measured NNDs to a simulated pattern of non-interacting molecules, modelled as complete spatial randomness (CSR). These CSR coordinates were simulated for each imaged cell within the binary mask and preserve measured protein densities (Methods and **Fig. S4**). For the K2 → ILK direction (**Fig. 5g** left), the CSR distribution for K2-LL/AA cells shifted towards larger distances compared to CSR distribution for K2-WT, directly reflecting the lower ILK density in K2-LL/AA FAs. Conversely, the CSR references for the ILK → K2 (**Fig. 5g** right) direction were indistinguishable across both cell lines, consistent with the constant K2 density.

A deviation of experimental NND distributions from their respective density-matched CSR baseline is indicative of specific interactions causing additional molecular proximity beyond random colocalization. In K2-WT cells, the average experimental NND distributions showed a significant enrichment of short distances as compared to the CSR reference for both NND directions, demonstrating molecular interactions that bring K2-WT and ILK into close proximity. In contrast, K2-LL/AA cells exhibited an average NND distribution from K2 to ILK consistent with CSR. While the reciprocal ILK → K2-LL/AA distribution approached its CSR control, it remained significantly distinct, indicating that the visually observed pairs of ILK and K2-LL/AA in **Fig. 5e** are not purely density-based random colocalization.

We next characterized this directional asymmetry between NND directions by calculating the percentage of K2 molecules with at least one ILK neighbour within 30 nm, alongside the reciprocal percentage of ILK molecules with a nearby K2 (**Fig. 5h**). While no discrete distance threshold can unambiguously distinguish an interacting from a non-interacting molecule based on proximity, even in the absence of labelling efficiency and linkage errors, this metric allows us to assess the relative likelihood of molecular association by comparison to the CSR baseline. In K2-WT cells, both directional percentages are enhanced over CSR. The percentage of K2-LL/AA molecules with a nearby ILK dropped toward CSR level, yet remained statistically distinct. While this residual enrichment was not captured in the average K2 → ILK NND distribution (**Fig. 5g**, bottom left), likely due to cell-to-cell variability, a small enrichment of short distances is visually apparent. This was corroborated by a second clone (clone 11, **Fig. S6a**,**b**) and NND curves for individual cells (**Fig. S7**), both of which revealed a small but significant degree of molecular proximity between K2-LL/AA and ILK that exceeds random colocalization.

The directional asymmetry becomes apparent in the percentage of ILK molecules that have a K2 within 30 nm (**Fig. 5h**). While the percentage of ILK with a nearby K2-LL/AA decreased compared to K2-WT, it remained substantially above the CSR baseline. In contrast, the reciprocal value of K2-LL/AA with nearby ILK dropped much closer to CSR. This difference indicates that although the majority of K2-LL/AA molecules are recruited to FAs without an ILK partner, a small pool of ILK molecules successfully localized to FAs maintains in a non-random proximity to K2-LL/AA above the density-based chance prediction. This observation was corroborated by analysis of individual cells (**Fig. S8**) of clone 11 (**Fig. S6c**).

Altogether these single-molecule measurements demonstrate that the K2-LL/AA impairs the FA recruitment as well as the nanoscale proximity to ILK, which identifies a direct interaction of K2-WT and ILK as the major recruitment mechanism. A fraction of ILK still successfully localizes to FAs and maintains a non-random spatial association with K2-LL/A. Although ILK may have residual direct or indirect interactions with K2-LL/A, the primary recruitment pathway of ILK to FAs is fundamentally compromised.

### K2-dependent recruitment of ILK to activated β1 Integrins

Finally, we sought to determine the recruitment mechanism of ILK to β1 integrin by assessing whether ILK, K2-WT and K2-LL/AA are recruited individually or together to activated β1 integrins. In three-target Exchange-PAINT, mCherry-ILK-WT, mEGFP-K2-WT or mEGFP-K2-LL/AA were labelled as described above, and activated β1 integrins were identified using the 9EG7 antibody with DNA-conjugated secondary FAB fragments (**Fig. 6a**). Protein maps were obtained from individual channels (**Fig. 6b–d**). Ternary complexes of active β1 integrin, ILK and K2-WT were readily observed (**Fig. 6d** top, regions **i** and **ii**), whereas active β1 integrin, ILK and K2-LL/AA were much less abundant (**Fig. 6d** bottom, regions **i** and **ii**). Binary complexes of active β1 integrin with either K2 (**Fig. 6d** top and bottom, region **iii**) or ILK (**Fig. 6d** top and bottom, region **iv**) were present in both K2-WT and K2-LL/AA expressing cells.

**Figure 6.**
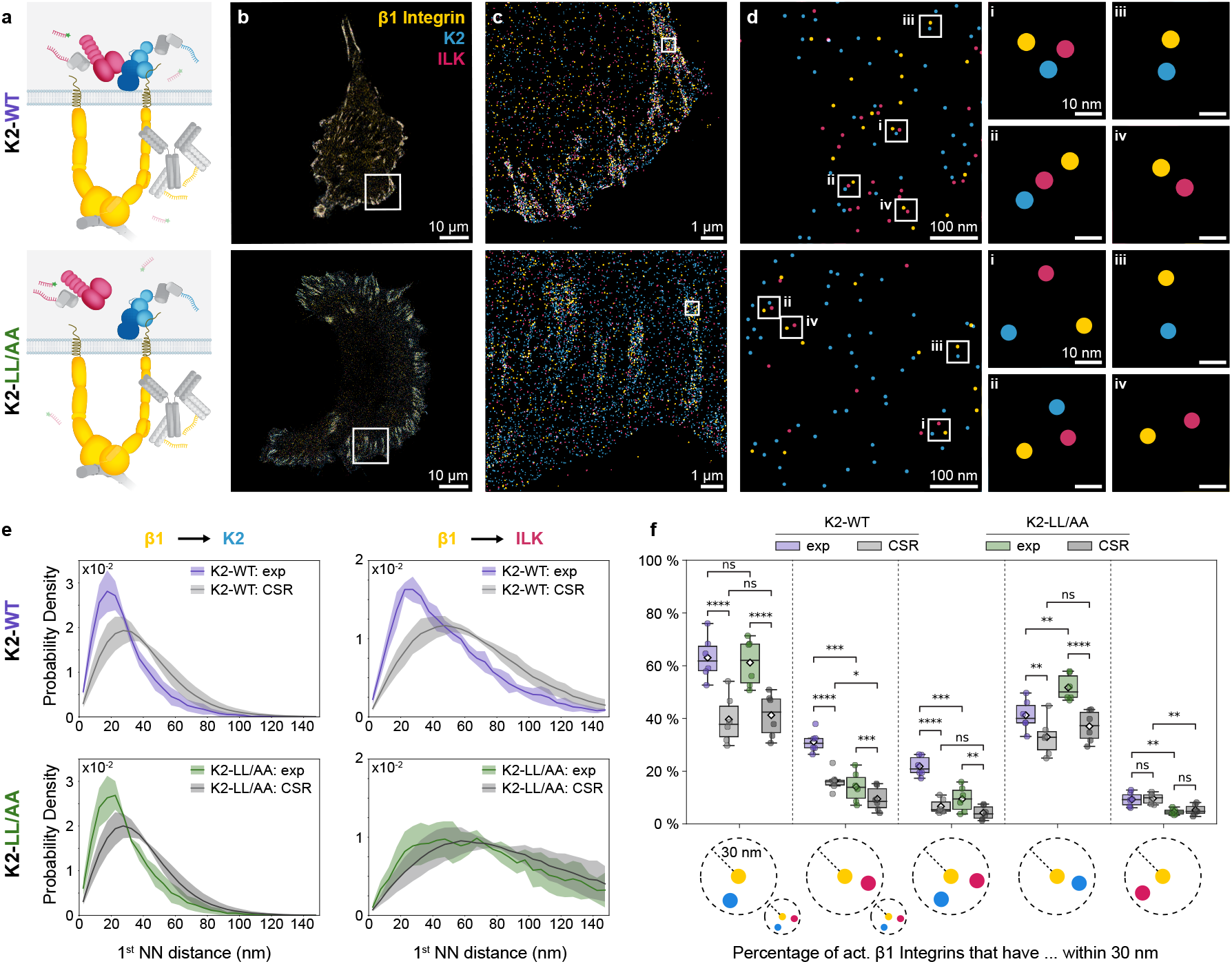
ILK recruitment to β1 integrins depends on K2. **a** Labelling strategy for three-plex Exchange-PAINT of mCherry-ILK, mEGFP-K2 (WT or LL/AA) and active β1 integrins. The latter are labelled via 9EG7 antibody and secondary DNA-coupled FAB fragments. **b** Representative three-plex Exchange-PAINT images of a K2-WT cell (top) and a K2-LL/AA cell (bottom). **c** Enlarged view of FAs. **d** Protein coordinates reveal ternary complexes of active β1 integrin, ILK and K2 (WT and LL/AA) (e.g. regions i and ii) and binary complexes of active β1 integrins and K2 (e.g. regions iii) as well as complexes of active β1-integrins and ILK (e.g. regions iv). **e** Average NND distributions from active β1 integrin to K2 or ILK are compared to simulated CSR scenarios (Mean ± SD of N = 6 cells per cell line). Left (β1 → K2): K2-WT and K2-LL/AA distributions are statistically indistinguishable (p = 0.995 for experimental and p = 0.999 for CSR distributions). Both are significantly closer to active β1 integrins than in absence of molecular interaction (p < 0.0001 respectively). Right (β1 → ILK): The experimental and CSR distributions in K2-LL/AA cells are shifted to larger distances compared to K2-WT (p < 0.0001 respectively). ILK is significantly closer to active β1 integrins only in K2-WT cells (p < 0.0001), but not in K2-LL/AA cells (p = 0.124). Modified Chi-squared test(56) at α = 0.05. **f** Percentage of active β1 integrins that have a K2 molecule, an ILK molecule, K2 and ILK simultaneously or only one of them within a radius of 30 nm are calculated. The threshold of 30 nm accommodates the linkage error of primary antibody / secondary Fab labelling for β1 integrin (∼ 15-20 nm) and nanobodies for mCherry-ILK and mEGFP-K2 (∼ 5-10 nm), alongside an average localization precision of ∼ 3 nm. Data from N = 6 cells per cell line. Boxplots show median and 25th and 75th percentile with whiskers reaching the last data point within 1.5 × interquartile range, diamonds indicate mean. Paired two-sample t-test for exp vs. CSR and Welch’s t-test for K2-WT vs. K2-LL/AA: ns (not significant) p > 0.05, * 0.01 < p ≤ 0.05, ** 0.001 < p ≤ 0.01, *** 0.0001 < p ≤ 0.001, **** p ≤ 0.0001.

To evaluate molecular interactions underlying these protein complexes, we performed CSR simulations to obtain a reference for the experimentally observed patterns. We first examined pairwise interactions by measuring the distance from each active β1 integrin to its nearest K2 molecule (**Fig. 6e** left, **Fig. S9a**) and found no difference between the distributions measured for K2-WT and K2-LL/AA, respectively. The K2-WT and K2-LL/AA localized significantly closer to β1 integrin than expected from random colocalization, which indicates that β1 tail recruitment is not influenced by ILK binding. When co-expressed with K2-WT, ILK localized significantly closer to active β1 integrins than in the CSR simulation (**Fig. 6e** right), indicating the involvement of a molecular interaction that brings ILK to β1 tails in FAs. In contrast, co-expression with K2-LL/AA resulted in an experimental NND distribution from β1 to ILK that overlaps with the CSR simulation. Thus, ILK was no longer recruited to β1 tails, owing to the impaired binding of K2-LL/AA to ILK. Comparing the wild type and mutant cases clearly indicates that ILK is indirectly recruited to β1 integrins through K2.

Although the NND curves from β1 to ILK in K2-LL/AA cells are not statistically different from CSR, the measured curve shows a slight enrichment of short distances compared to CSR. This is also apparent in single-cell NND curves (**Fig. S9b**) and in clone 11 cells (**Fig. S10a**). These findings strongly suggest that ILK recruitment to active β1 integrins is primarily mediated by binding to K2-WT, and that residual recruitment may stem from other proteins that bridge ILK and active β1 integrins, either with or without the involvement of K2. Further possibilities are weak direct interactions between ILK and β1 tails or residual binding of ILK to K2-LL/AA.

To investigate these possibilities, we quantified the exclusive and simultaneous occurrence of K2 and ILK within 30 nm of active β1 integrins (**Fig. 6f**). Ternary complexes of active β1, ILK and K2-WT were significantly enriched relative to CSR. In contrast, ternary complexes containing K2-LL/AA were slightly more frequent than CSR. This recapitulates that most ILK proteins are recruited to β1 via the K2 bridge. Next, we measured the exclusive localization of K2 at active β1 and found that both K2-WT and K2-LL/AA levels exceeded CSR levels, confirming that ILK is not required for K2 binding to β1-tails. Conversely, colocalization of ILK at active β1 in the absence of K2 was reflected random colocalization. These observations were confirmed in clone 11 cells (**Fig. S10b**) and single-cell analysis (**Fig. S11**), and indicate that neither direct ILK-β1 tail binding nor an alternative, K2-independent bridging molecule mediate β1-tail recruitment of ILK in K2-LL/AA expressing cells. We conclude that K2 is an obligatory component in positioning ILK in the vicinity of β1. However, the possibility of a larger multi-protein complex where K2 and another adapter protein cooperate to crosslink β1 and ILK cannot be excluded. The slight enrichment of β1, ILK and K2-LL/AA complexes above CSR possibly reflects either such cooperative interactions or residual binding of K2-LL/AA to ILK. Altogether, our findings demonstrate that ILK cannot directly bind β1 tails in cells, and that ILK is entirely recruited by K2-dependent mechanisms to active β1 integrins, predominantly via K2 directly connecting with ILK and β1.

## Discussion

The recruitment of ILK to the cytoplasmic domain of β1 integrin remains a topic of debate within the integrin community. ILK was initially identified as an integrin tail-interacting protein using genetic and biochemical binding assays. Subsequent studies refuted this finding and proposed an indirect binding to integrin tails. We investigated this possibility using an interdisciplinary approach, and report that K2 serves as the primary platform to recruit ILK and the interacting proteins PINCH and parvin to β1 tails in FAs of fibronectin-seeded fibroblasts.

Our findings, supported by a series of *in vitro* experiments combined with *in situ* super-resolution microscopy, demonstrate that the full-length IPP complex does not directly bind to integrin cytoplasmic tails. The K2-LL/AA mutation, which was abrogates kindlin-ILK interaction, significantly reduced ILK localization to active β1 integrins in cells, providing strong support for the notion that ILK recruitment is an indirect event mediated by kindlin. This raises the question of whether ILK is recruited as part of the intact IPP complex or whether its components can act independently at integrin adhesion sites.

The stability of the IPP complex in mammalian cells is well-established(47, 48), and our biochemical data are consistent with this. However, evidence from Drosophila suggests that ILK can be recruited to integrin adhesion sites independently of PINCH(49). Furthermore, post-translational modifications and signalling events in cultured mammalian cells may lead to partial uncoupling of IPP components followed by context-dependent functions of ILK, PINCH, and parvin outside of the canonical trimer(48, 50).

While we cannot exclude the possibility that isolated ILK binds directly to integrin cytoplasmic tails, our cellular analyses clearly show that recruitment of ILK and α-parvin to integrins is kindlin-dependent.

We also observed that K2 recruitment to FAs and its binding to integrins occur independently of ILK binding. The ILK-binding-deficient K2-LL/AA protein localized to FAs and in close proximity to β1 integrins. This finding contrasts with previous studies showing that ILK influences K2 localization. These studies examined kindlin mutations in the presence of endogenous kindlin(16, 17, 19), which likely outcompeted the mutant protein during FA recruitment. In *C. elegans*, ILK binding induces conformational changes in kindlin(13) that are important for subsequent integrin association. However, our results along with other research with mammalian cells demonstrate species-specific differences in the interaction. In mammals, mutations like S351V in K2, which align with critical residues in *C. elegans*, do not impair ILK binding or K2 localization(17).

Our *in situ* microscopy reveals a two-step recruitment mechanism for the IPP complex to FAs: recruitment to the β1-tails is strictly K2 dependent, whereas the localization to FAs is dependent on K2/ILK interaction, but might also involve further, probably minor, K2-independent mechanisms. This is in line with studies describing distinct pools of FA proteins – dynamic and stably associating pools – depending on the FA’s location and orientation relative to the cell edge(51), which supports the idea that protein recruitment and dynamics within FAs are influenced by local environmental factors and specific PPIs. Despite our evidence for K2-dependent ILK recruitment, we observed residual ILK-integrin proximity in cells expressing K2-LL/AA. This could result from weak residual binding between ILK and the K2-LL/AA mutant, although NMR experiments demonstrated that the K2-LL/AA substitutions abolish ILK binding(16). Alternatively, it is possible that K2 positions ILK near integrin tails via another K2 and ILK binding partner. This scenario of a multi-protein assembly involving at least four components (β1, K2, an intermediate protein, and ILK) remains consistent with our observation that K2 is strictly required for ILK localization near β1 tails. The identity of the kindlin-ILK bridging molecule is unknown. Our experiments exclude a role for paxillin. However, we can neither exclude other paxillin family members like leupaxin and Hic-5 nor other FA proteins.

Our workflow successfully disentangles the stepwise recruitment process of ILK to FAs and β1-integrin tails, a mechanism that is obscured at diffraction-limited length scales and highlights the complex and spatially organized nature of FAs. While our measurements report proximity relationships rather than fully resolved nanoscale domains, they are consistent with a view in which FAs are not randomly organized but instead exhibit both vertical layering(52) and lateral heterogeneity(53-55) in protein composition.

The combination of *in vitro* assays with *in situ* microscopy at single molecule resolution allows for the extraction of spatial protein patterns with highest sensitivity. By comparing patterns of wild-type and mutant cells to reference simulations, we translate proximity metrics into molecular interaction networks with high confidence. *In vitro* assays directly measure interactions, but only in isolated and potentially artificial settings. While the demonstrations of *in vitro* interactions are an essential first step in PPI analysis, these interaction patterns may not accurately reflect those in a natural cellular context. Exchange-PAINT provides single-cell molecular protein maps that reveal spatial patterns from nanoscale protein colocalization to their organization into larger structures such as FAs. Furthermore, by comparing proximity metrics derived from experimental and simulated CSR protein maps, we effectively distinguish molecular interactions from random colocalization. Importantly, our approach used protein densities and FA masks extracted from measured molecular maps as input parameters for CSR simulations. This ensures direct comparability between *in situ* measurements and *in silico* data, providing robust validation of observed interactions. Furthermore, by comparing wild-type and ILK-binding-deficient K2 in cells, our pipeline distinguishes whether observed interactions are direct or mediated by additional molecules, thereby providing robust *in situ* protein-protein interaction (PPI) data. In this way, our workflow offers mechanistic insights into ILK recruitment to β1 integrins in cells and therefore represents the final proof of the *in vitro* biochemical characterization: we identified the kindlin-dependent recruitment of ILK to β1-integrin as major pathway that operates in cells.

## Acknowledgments

We thank Sebastian Strauss and Jisoo Kwon for binder conjugation and Thomas Schlichthärle for initial experiments. The authors thank the Core Facilities at the MPI of Biochemistry for their support. Specifically, the Protein Production Facility for expression of the IPP complex and kindlin proteins, the Biochemistry Core Facility for peptide synthesis and ITC measurements, and the CryoEM Facility for the use of the mass photometry. S.C.M.R. acknowledges the support by the IMPRS-ML graduate school.

## Author Contributions

S.C.M.R. performed Exchange-PAINT experiments, developed the analysis software and analyzed Exchange-PAINT and confocal microscopy data. R.T.B. generated and characterized the cell lines, performed the integrin tail ELISA experiments, and conducted confocal microscopy imaging. F.B. purified recombinant proteins and performed biophysical characterizations. J.D.S. performed kindlin2-ILK immunoprecipitations. S.C.M.R, R.T.B., R.J. and R.F. interpreted data and wrote the manuscript. R.J. and R.F. conceived and supervised the study. S.C.M.R. and R.T.B. contributed equally. All authors reviewed and approved the final manuscript.

## Data availability

The single-molecule localization data generated in this study is available via Zenodo at https://doi.org/10.5281/zenodo.19921201.

## Code availability

DNA-PAINT image processing can be performed using Picasso available via GitHub at https://github.com/jungmannlab/picasso. The custom-written scripts used in this study are available via GitHub at https://github.com/jungmannlab/adhesion_ppi.

