## Supplementary Information for "ILK binding to β1 Integrin is indirectly mediated by Kindlin-2"

**This PDF file includes:**

Supporting Information Text
Figures S1 to S12
SI References

**Supporting Information Text**

**Materials.** DNA oligonucleotides modified with C3-azide and Cy3B were purchased from Metabion and MWG Eurofins. Fetal Bovine Serum FBS (10500-064), 1x PBS (pH 7.2; 20012-019), EDTA 0.5 M (pH 8.0; AM9260G), 0.05% trypsin–EDTA (25300-054), Salmon Sperm DNA (15632011), sodium chloride 5 M (cat: AM9759), ultrapure water (cat: 10977-035) were purchased from Thermo Fisher Scientific. Sodium hydroxide (cat: 31627.290) was purchased from VWR. Tween-20 (P9416-50ML), (±)-6-hydroxy-2,5,7,8-tetra-methylchromane-2-carboxylic acid (trolox; 238813-5G), Sodium azide (769320), Bovine serum albumin BSA (A4503-10G) and methanol (cat: 32213-2.5L) were ordered from Sigma-Aldrich. Paraformaldehyde (15710) were obtained from Electron Microscopy Sciences. Triton X-100 (6683.1) was purchased from Carl Roth. 90-nm gold nanoparticles (G-90-100) were ordered from Cytodiagnostics.

**Antibodies.** The following antibodies or molecular probes were used at indicated concentrations for western blot (WB) or immunofluorescence (IF): anti-kindlin-2 (MAB2617; Millipore) WB 1:1,000; anti-ILK (#611803; BD Bioscience) WB: 1:2,000; anti-talin (8D4; Sigma) W: 1:1,000; anti-β-actin (A5441; Sigma-Aldrich) WB 1:2,000; anti-GFP (11814460001; Sigma-Aldrich) WB 1:1000; anti-α-parvin (#8190; Cell Signaling) IF: 1:500; anti-paxillin (#610051; BD Bioscience) IF 1:500, anti-vinculin (V9131, Sigma-Adrich) IF 1:500.
The following secondary antibodies were used: goat anti-mouse Alexa 546 (A11003), donkey anti-rabbit Alexa 546 (A10040), goat anti-rabbit Alexa 647 (A21244) (all from Invitrogen) IF: 1:500; donkey anti-rabbit Cy3 (711-165-152) (from Jakson ImmunoResearch) IF: 1:500; goat anti-mouse HRP (172-1011) and goat anti-rabbit HRP (172-1019) (both from Biorad) W: 1:10,000. Phalloidin-Alexa 488 (A12379; Thermo Fisher Scientific; 1:500) and DAPI (Sigma) were used to stain F-actin and nuclei, respectively.

**Constructs and plasmids.** A vector expressing mEGFP-tagged K2 was generated by replacing the EGFP sequence in the pEGFP-C1-K2 plasmid(1) with mEGFP. ILK cDNA was cloned into the pmCherry vector (Clontech). Point mutations into the mouse K2 (L353A/L357A) and ILK (K423D/I427E) cDNAs were introduced by site-directed mutagenesis using PfuUltra II (600670, Agilent Technologies) according to the manufacturer's protocol. For stably expressing the K2 and ILK cDNAs, the cDNAs were cloned into the retroviral expression vector pRetroQ. For CRISPR-mediated depletion of mouse Paxillin and mouse ILK the guide-RNAs were ligated into pSpCas9(BB)-2A-Puro vector V2.0 (gift from Feng Zhang (Addgene plasmid # 62988; http://n2t.net/addgene:62988; RRID:Addgene_62988)(2). The mouse ILK cDNA carrying two point mutations (C346S; C422S) to increase the protein solubility of the complex(3), PINCH-1 cDNA and N-terminally His tagged β-parvin cDNA were cloned into the pACEBac1 baculovirus transfer vector that serves as an acceptor vector in the MultiBAC system (Geneva Biotech). The correct sequences of all constructs were verified by Sanger sequencing (Eurofins).

**Stable viral transduction.** To generate stable cell lines, HEK293T cells were transiently transfected with vesicular stomatitis virus G glycoprotein (VSV-G) pseudotyped retroviral vectors. Viral packaging plasmids were transfected, and viral particles were harvested by collecting culture supernatant at 48 and 72 h post-transfection. The supernatant was filtered through 0.45 μm membranes, and viral particles were pelleted by ultracentrifugation at 20,300 rpm for 2 h with a SW 32 Ti rotor (Beckman Coulter). Viral pellets from each 15-cm dish were resuspended in 45 μl cold Hank's balanced salt solution (Thermo Fisher Scientific, 14175046). For infection, 5-10 μl of viral suspension was added to 60,000 cells seeded in 6-well plates the day before. Following infection, cells were sorted to distinct expression levels by fluorescence-activated cell sorting on a BD FACSAria III (BD Biosciences).

**Cell lines.** HEK293T cells (ATCC, CRL-3216), SV40 large T-immortalized mouse kidney fibroblasts lacking talin-1, talin-2, kindlin-1, and kindlin-2 (qKO cells), as well as the parental Flox cells(1) were cultured in Dulbecco's modified Eagle's medium (DMEM; Gibco) with 10% fetal bovine serum (FBS; v/v; Gibco) and 1% penicillin–streptomycin (P/S; v/v; Gibco) at 37 °C, 5% CO2 and

95% humidity. To generate qKO Paxillin and ILK knock-out (KO) lines, qKO cells were transiently transfected with CRISPR-hSpCas9 vectors encoding the specific guide-RNAs. Following 48 h selection with 2 µg/ml puromycin, single-cell clones were isolated by FACS (BD FACSAria III (BD Biosciences)). KO clones were confirmed by WB analysis. For rescue experiments, qKO/ILK-KO cells were reconstituted with mouse talin-1 and combinations of N-terminally mCherry-tagged ILK (wild-type mCherry-ILK (ILK-WT) or kindlin-binding deficient mCherry-ILK K423D/I427E (ILK-KI/DE), together with N-terminally mEGFP-tagged K2 (wild-type mEGFP-K2 (K2-WT) or ILK-binding deficient K2 (mEGFP-K2 L353A/L357A (K2-LL/AA)). qKO/Pax-KO clones were reconstituted with mouse talin-1 and either mEGFP-K2-WT or mEGFP-K2-LL/AA, supplemented along with mCherry-tagged paxillin. All cell lines were routinely tested for mycoplasma contamination.

**Protein expression and purification. Kindlin-2.** K2, K2-QW/AA, and K2-LL/AA were expressed and purified as described earlier (Böttcher et al.(1)). Proteins were expressed in E. coli Rosetta (Millipore) following induction with 0.2 mM IPTG at 18 °C overnight. Cells were lysed, and proteins were purified by immobilized metal chelate affinity chromatography (IMAC) in high-salt TBS buffer (20 mM Tris, pH 7.5, 500 mM NaCl, 1 mM Tris(2-carboxyethyl)phosphine (TCEP)). The $His_{10}$–SUMO tag was cleaved overnight with SenP2 protease (obtained from the Max Planck Institute of biochemistry (MPIB) Core Facility), and the cleaved protein was further purified by size exclusion chromatography (Superdex 200 16/60; GE Healthcare) using TBS (20 mM Tris, pH 7.5, 200 mM NaCl, 1 mM TCEP) supplemented with 5% glycerol. **IPP complex.** IPP complex was produced in Sf21 cells using the MultiBac system. Cells from 500 ml liquid culture were collected and lysed in 40 ml lysis buffer (50 mM HEPES, pH 7.6, 150 mM NaCl, 10 µM ZnCl2, 0.1% Tween-20, 1 mM PMSF, 1 mM TCEP, 40 µL DNAse (obtained from the MPIB Core Facility) for 5 ml cell pellet, 0.1 mM ATP and 5 mM MgCl2). Lysis was performed on ice using a Dounce tissue grinder. After incubation for 10 min on ice, the lysate was cleared by centrifugation at 58,000 x g for 60 min at 4°C. The supernatant was filtered through 0.45 µm syringe filter, and imidazole was added to a final concentration of 40 mM. The tagged complex was purified by immobilized metal affinity chromatography using a 5 ml HisTrap HP column (GE Healthcare, 17-5248-02) on a BioRad NGC chromatography system using Buffer A (50 mM HEPES, pH 7.5, 150 mM NaCl, 10 µM ZnCl2, 0.05% Tween-20, 1 mM TCEP, 0.1 mM ATP and 5 mM MgCl2) and Buffer B (50 mM HEPES, pH 7.5, 150 mM NaCl, 500 mM imidazole, 10 µM ZnCl2, 0.05% Tween-20, 1 mM TCEP, 0.1 mM ATP and 5 mM MgCl2). The column was washed with 15% Buffer B, and the bound complex was eluted with 40 % Buffer B. The His-tag was removed by cleavage with PreScission Protease (2 µl per 100µg protein) for 1.5 h on ice. The sample was subsequently purified by SEC using a Superdex 200 pg column (Cytiva) equilibrated in Buffer C (50 mM Citrate, pH 6.4, 150 mM NaCl, 1 mM TCEP, 0.1 mM ATP and 5 mM MgCl2). Purified complex was snap-frozen in liquid nitrogen and stored at -80°C.

**Synthetic integrin cytosolic peptides.** Synthetic integrin cytosolic peptides were synthesized in house by the Max Planck Institute of Biochemistry Core Facility. The correct peptide sequence was controlled by high-resolution intact mass spectrometry. Mouse β1 wt cytoplasmic tail peptide (HDRREFAKFEKEKMNAKWDTGENPIYKSAVTTVVNPKYEGK-OH),
β1 scrambled peptide (EYEFEPDKVDTGAKGTKMAKNEKKFRNYTVHNIWESRKVAP-OH), β1-YY/AA peptide (HDRREFAKFEKEKMNAKWDTGENPIAKSAVTTVVNPKAEGK-OH)
mouse β3 wt cytoplasmic tail peptide (HDRKEFAKFEEERARAKWDTANNPLYKEATSTFTNITYRGT-OH), β3 scrambled peptide (RRIESFNAGKTEEDRANTYWLAFPEETKYRAHKTTDTFNAK-OH)

**Integrin peptide ELISA.** Synthetic integrin cytosolic peptides (β1-WT, β1-YY/AA, β1-Scr, β3-WT, and β3-Scr) were dissolved in PBS to a final concentration of 500 nM. Aliquots of 100 µl were immobilized on 96-well MaxiSorp plates (Nalge Nunc) overnight at 4°C. Plates were blocked with 200 µl PBS containing 3% BSA for 1 h at room temperature and then washed thrice with 200 µl PBS. Different dilutions of recombinant IPP and kindlin proteins were prepared in PBS supplemented with 3% BSA (100 µl/well). Plates were incubated overnight at 4°C and then washed three times with 200 µl Mammalian Protein Extraction Reagent (M-PER buffer, Thermo Fisher

Scientific, 78501). Detection was performed by adding 100 µl of anti-ILK or anti-kindlin-2 antibody (1:1,000 in PBS plus 3% BSA), followed by overnight incubation at 4°C. After washing thrice with 200 µl MPER buffer, wells were incubated with anti-rabbit-IgG HRP-conjugate or anti-mouse-IgG HRP-conjugate (1:10,000 in PBS supplemented with 3% BSA) for 1 h at room temperature. After the final three washes with MPER buffer (200 µl/well) PBS, and one PBS washing step, the assay was developed with 100 µl of ABTS substrate solution (1-Step ABTS; Thermo Fisher Scientific). Absorbance was measured absorbance at λ = 405 nm in a SpectraMax ABS Plus plate reader (Molecular Devices).

**Spreading and adhesion assays.** Cells were grown to 70% confluence followed by overnight incubation in DMEM containing 0.2% fetal calf serum (FCS). Following detachment with trypsin/EDTA, cells were serum-starved for 1 h at 37°C in adhesion assay medium (10 mM HEPES pH 7.4; 137 mM NaCl; 1 mM $MgCl_2$; 1 mM $CaCl_2$; 2.7 mM KCl; 4.5 g/l Glucose; 3% BSA (w/v)) or DMEM containing 3% BSA.
For adhesion assays, flat-bottom 96-well plates were coated with increasing concentrations of FN (0.3 – 9.0 µg/ml Calbiochem) and subsequently blocked for 1 h with PBS containing 3% BSA. A total of 40,000 cells per well were plated in adhesion buffer or DMEM containing 3% BSA and incubated for 30 min at 37°C. Non-adherent cells were removed by washing with PBS, and adherent cells were fixed with PBS containing 4% PFA. Adherent cells were stained with crystal violet (0.1% in 20% methanol) for 15 min, washed extensively with H2O, and the dye was released using 2% SDS/H2O. Absorption was measured at λ=570 nm in a SpectraMax ABS Plus plate reader (Molecular Devices).
For cell spreading, 40,000 cells were seeded on 10 µg/ml FN-coated coverslips, cultured in DMEM containing 3% BSA at 37°C, fixed with 4% PFA (w/v) in PBS and stained with phalloidin-Alexa488 and DAPI. At least 30 cells from six randomly selected fields per condition were imaged using a Zeiss LSM 780 confocal laser scanning microscope equipped with 40x/NA 1.4 oil immersion objective and cell spreading area was quantified using ImageJ (release 1.53, (Schneider et al., 2012)).

**GFP-immunoprecipitation.** Cells were lysed in M-PER buffer supplemented complemented with 5% protease inhibitor cocktail (Roche; 04693159001), sonicated, and centrifuged at 14,000 rpm for 10 min at 4 °C. For GFP-IPs, 20 µl GFP-Trap Magnetic Agarose beads (Chromotek; GTMA-20) were washed 3x with M-PER buffer and incubated with 1.2 mg lysate for of 2 h at 4 °C. After three washes with M-PER buffer, bound proteins were eluted by adding 60 µl of 4× Laemmli sample buffer (500 mM Tris-HCl, pH 6.8; 8.0% SDS; 40% glycerol; 20 mM EDTA; 0.03% bromophenol blue) supplemented with 0.33 g urea per ml, and the bead suspension was heat-denatured at 95 °C for 6 min.

**Mass photometry.** Mass photometry measurements were carried out in silicone gaskets on microscopy slides that had been cleaned by consecutive sonication in Milli-Q water, isopropanol, and Milli-Q water. Mixtures of freshly prepared, purified proteins (typical concentration of proteins were 5–25 nM) were added to the gaskets containing 4 µl buffer and images were acquired for 60 s at a OneMP mass photometer (Refeyn Ltd, Oxford, UK). Data acquisition was performed using AcquireMP (Refeyn Ltd) and all images were processed and analyzed using DiscoverMP (Refeyn Ltd, v1.2.3). Sizes were calculated by standardizing measurement results against a size standard (NativeMark™ Unstained Protein Standard, Thermo Fisher).

**Isothermal titration calorimetry (ITC).** Quantitative ITC-measurements were run on a PeaqITC instrument (Malvern) at a constant jacket temperature of 14°C. IPP complex and kinlin-2 were rebuffered in buffer A (50 mM Citrate, pH 6.4, 150 mM NaCl, 1 mM TCEP, 0.1 mM ATP and 5 mM $MgCl_2$) and β1 and β3 tail peptides were dissolved in and dialysed against excess volumes of buffer A to remove all potential contaminants and guarantee good buffer matching to the IPP complex. Twelve to eighteen times 3 µl of the respective β-tail peptide were injected into the measurement cell containing IPP or K2. All data were analysed using MicroCal PeaqITC Analysis Software.

**FA Intensity and Morphology Analysis for Confocal Data.** Three-channel confocal data was analysed in Fiji(4) using custom Macros. Cells were segmented via the Kindlin channel. We used the default thresholding function with a minimum intensity of 2500, converted the thresholded image to a mask and applied a Gaussian blur (radius = 4 pixel). After another conversion to a binary image, we identified the background area as the black regions in the mask. Furthermore, we used the Analyze Particles function with a minimum particle size of 10000 pixels to identify cell outlines in the mask. Segmentation results, the background area and cell outlines, were saved as ROIs. Successful segmentation was verified for each FOV. If cells were not properly segmented, ROIs were corrected manually. Subsequently FA areas were identified using paxillin or Vinculin (in Pax-KO cells) as FA markers. For each cell individually, automatic thresholding was applied. Conversion to a mask, Gaussian blur (radius = 1 pixel) and another binary conversion resulted in the final FA ROIs. These were again manually corrected if the automatic thresholding clearly did not identify the FAs correctly. Mean intensities of K2 and α-parvin were measured within every cell's FA ROI and within the background ROI. Using python, we performed background correction for each FOV by subtracting the respective mean background intensity from each cell's intensity value. Normalized fluorescent intensities were calculated as a ratio of the target protein signal to that of the FA marker (paxillin and vinculin) or K2. Binary masks were created from cell and FA ROIs. Cell and FA areas as well as FA aspect ratio were extracted using the Analyze Particles function. To quantify FA spatial distribution relative to the cell edge, a Euclidian distance map was generated from each cell's boundary ROI. FA centroids were extracted from Analyze Particles and the distance of a FA centroid to the cell edge was determined as the exact pixel value of the centroid on the distance map. Cells touching the edge of the field of view were excluded for counting the number of FAs per cell, cell spreading area and total adhesion area, but were included in intensity measurements and metrics referring to individual adhesions.

**Buffers for DNA-PAINT Imaging**

- Blocking buffer: 1xPBS, 1mM EDTA, 0.02% Tween-20, 0.05% NaN3, 2% BSA, 0.05 mg/ml sheared salmon sperm DNA
- 100x Trolox was made by adding 100 mg Trolox to 430 µl of 100% methanol and 345 µl of 1 M NaOH in 3.2 ml ultrapure water.
- Buffer C: 1× PBS, 1 mM EDTA and 500 mM NaCl, pH 7.4; 0.02% Tween-20
- Imager solution: Buffer C, 1x Trolox, supplemented with Cy3B-coupled DNA imager strands.

**Nanobody-DNA conjugation.** Nanobodies against GFP (clone 1H1, Cat No: N0305 and 1B2, custom order) and mCherry (clone 2B12, Cat No: N0405) were purchased from NanoTag Biotechnologies with a single ectopic cysteine at the C-terminus for site specific and quantitative conjugation. Nanobodies were conjugated as described previously(5). First, buffer was exchanged to 1× PBS + 5 mM EDTA, pH 7.0 using Amicon centrifugal filters (10k MWCO). Free cysteines were reacted with 20-fold molar excess of bifunctional Sulfo DBCO-PEG4-Maleimide linker (BroadPharm, cat: BP-23318) for 2-3 hours on ice. Unreacted linker was removed by buffer exchange to PBS using Amicon centrifugal filters. Azide-functionalized DNA (5xR1 for anti-mCherry nanobody: TCCTCCTCCTCCTCCTCCT; 7xR3 for anti-GFP nanobodies: CTCTCTCTCTCTCTCTCTC) was added with 5-10 molar excess to the DBCO-nanobody and reacted overnight at 4°C. Unconjugated nanobody and free azide-DNA was removed by anion exchange using an ÄKTA Pure liquid chromatography system equipped with a Resource Q 1-ml column.

**Fab-DNA conjugation.** Fab Fragment Donkey Anti-Rat (Jackson ImmunoResearch, cat: 712-007-003) was conjugated to azide-functionalized DNA using DBCO-sulfo-NHS ester crosslinker (Jena Bioscience GmbH, cat: CLK-A124-10). First, the FAB fragment was concentrated in 1xPBS with 30 kDa Amicon centrifugal filters and DBCO-sulfo-NHS crosslinker was added at 10× molar excess and incubated for 2 h at 4 °C on a shaker. To remove potential aggregates, the FAB fragments were centrifuged at 20000 g. Unreacted crosslinker was removed from the supernatant using Zeba spin desalting columns (7k MWCO) were used to remove unreacted crosslinker. Azide-DNA (2xR4: ACACACACA) was added to the purified FAB-crosslinker solution at 5× molar excess and incubated overnight at 4 °C. Subsequently, Amicon spin filters equilibrated in storage buffer (1xPBS pH 7.2 + 1 mM EDTA, pH 8 + 0.05% NaN3) were used to remove free azide-DNA and the FAB conjugate was stored at 4 °C.

**Super-resolution microscope setup.** Fluorescence imaging was carried out on an inverted microscope (Nikon Instruments, Eclipse Ti2) with the Perfect Focus System, applying an objective-type TIRF configuration equipped with an oil-immersion objective (Nikon Instruments, Apo SR TIRF×100, NA 1.49, Oil). A 560-nm laser (MPB Communications, 1 W) was used for excitation. The laser beam was passed through a cleanup filter (Chroma Technology, ZET561/10) and coupled into the microscope objective using a beam splitter (Chroma Technology, ZT561rdc). Fluorescence was spectrally filtered with an emission filter (Chroma Technology, ET600/50m and ET575lp) and imaged on an sCMOS camera (Hamamatsu, ORCA-Fusion BT or Andor, Zyla 4.2 Plus) without further magnification, resulting in an effective pixel size of 130 nm (after 2×2 binning). Scan speed was set to “standard” (Hamamatsu, ORCA-Fusion BT) or 540 MHz (Andor, Zyla 4.2 Plus). Images were acquired by choosing a square region of interest. Raw microscopy data was acquired using µManager(6) (Version 2.0.1).

**Cell culture and DNA-PAINT sample preparation.** Cells were maintained in high glucose Dulbecco’s modified Eagle’ medium (Thermo Fisher, 31966047) supplemented with 10% fetal bovine serum (Thermo Fisher, 10270106). For imaging, 10,000 - 20,000 cells were seeded on ibidi eight-well high glass-bottom chambers (no. 80807) per well with coating of 10 µg/ml fibronectin (Sigma Aldrich, cat: F4759). Cells were fixed with pre-warmed 4 % paraformaldehyde in PBS for 30 min at room temperature. After fixation, cells were washed three times with PBS. Gold nanoparticles (200 µl, diluted 1:2) were incubated for 5 min and washed three times with PBS. Blocking and permeabilization were performed with 0.2 % Triton X-100 in blocking buffer for 30 min. After washing with PBS, cells were incubated with 200 µl of purified rat anti-mouse CD29 clone 9EG7 antibody at 50 µg/ml (BD Pharmingen, cat: 553715) in blocking buffer overnight at 4 °C. Unbound antibodies were removed by washing three times with PBS, followed by washing once with buffer C for 10 min. 200 µl of blocking buffer containing 25 nM of secondary anti-rat FAB coupled to DNA sequence 2xR4, 25 nM of anti-GFP nanobody clone 1H1 and 25 nM of anti-GFP nanobody clone 1B2 both conjugated to 7xR3 and 25 nM of anti-mCherry nanobody clone 2B12 coupled to 5xR1 were incubated for 1 h at RT. Unbound probes were removed by washing three times with PBS, followed by washing once with buffer C for 10 min. Postfixation was performed with 4 % paraformaldehyde in PBS for 10 min before washing 3× with PBS.

**DNA-PAINT imaging.** The first Exchange-PAINT round targeting ILK was acquired using the imager solution supplement with R1 7nt imager (AGGAGGA-Cy3B). Between imaging rounds the sample was washed with 2–3 ml of PBS until no residual signal from the previous imager solution was detected. Then, the R3 7nt imager solution (GAGAGAG-Cy3B) was introduced and K2 was imaged. After a final washing step, active β1-integrin imaging was conducted using the R4 6nt imager solution (GTGTGT-Cy3B). ILK was imaged at imager concentrations between 200 and 400 pM, active β1 integrins at concentrations between 75 pM and 200 pM and K2 at 75 pM. Imaging was conducted at 63 mW at the objective at readout mode 2 (Hamamatsu, ORCA-Fusion BT) or a readout rate of 520 MHz (Andor, Zyla 4.2 Plus). In each round 50,000 frames were acquired at an exposure time of 75 ms. Active β1-integrin imaging rounds show a median NeNA precision of 2.95 nm [2.71 nm, 3.29 nm] (Median, 25th and 75th percentile), K2 rounds yield 2.78 nm [2.52 nm, 3.00 nm] and ILK rounds yield 3.09 nm [2.92 nm, 3.55 nm].

**Image analysis - DNA-PAINT image reconstruction.** Raw fluorescence data were subjected to super-resolution reconstruction using the Picasso software package(7) (latest version available at https://github.com/jungmannlab/picasso). Drift correction was performed with a redundant cross-correlation and gold particles as fiducials. Gold particles were also used to align all rounds for multiplexed Exchange-PAINT experiments. After channel alignment, localizations from gold particles and cells visible at the edge of the FOV were picked and removed in Picasso to perform the subsequent analysis only on the centred cell. DNA-PAINT localizations were analysed using the previously described(8) Picasso clustering algorithm (latest version available at https://github.com/jungmannlab/picasso) for each target individually. Circular clusters of localizations cantered around local maxima were identified and grouped (assigned a unique identification number). Localization groups were filtered to exclude clusters that originate from unspecific sticking of imagers to the sample by checking “Frame analysis” in Picasso. Subsequently, the canters of the localization groups were calculated as weighted mean by employing the squared inverse localization precisions as weights. The resulting centres represent individual protein coordinates. Overlaying protein coordinates of all rounds yields the final multiplexed DNA-PAINT image.

**Image analysis - Identification of FAs.** Kindlin signal was used as a marker to identify FAs as high-density regions. Thus, protein coordinates of the K2 channel were rendered as an image by creating a 2D histogram with 30 nm pixel size and applying a Gaussian blur of 90 or 120 nm. A binary mask is created by applying a custom threshold dividing the field of view (FOV) in high- and low-density regions, representing FAs and the rest of the FOV respectively. The mask is then smoothed to a final pixel size of 10 nm by spline interpolation. Values of 1 in the mask represent FAs, values of 0 represent the rest of the FOV. Finally, localizations inside of the FA mask are identified and kept for further analysis. Note that the mask based on K2 signal is used to filter out non-FA localizations of all three protein channels.

**Image analysis - Identification of protein complexes.** Pairwise interaction of act. β1, K2 or ILK was studied via nearest neighbour distances between their protein coordinates while considering only the proteins inside the FA mask. Ternary complex formation of all three species was assessed by defining a circle of 30 nm around every act. β1 protein. Then the percentage of act. β1 proteins, which have a certain combination of K2 and ILK within this radius, was calculated. All of these metrics are calculated based on protein coordinates identified in FAs of a single cell and thus limited in their precision by the finite number of proteins. By performing bootstrapping we generate 20 samples to estimate the range of possible outcomes given the same underlying distribution (shown in Supplementary Figures 7,8,9 and 11).

**Image analysis - Comparison of protein interactions to CSR simulations across cell lines and single cells**. The proximity between the three protein species and the amount of detected complexes that is expected in the absence of any direct molecular interaction is estimated via a simulated three-plex CSR (complete spatial randomness) dataset. For a given cell, comparability of the CSR distributions to the experimental distributions is ensured by using the measured protein densities as input for CSR simulation and simulating the protein patterns within the FA region of the mask. The latter ensures that boundary effects occurring e.g. in the calculation of nearest neighbour distances at the edges of FAs are identical in experimental and simulated data. By repeating the CSR simulation 20 times and proceeding with the mean of the respective output metric, fluctuations due to finite sample size are reduced. Supplementary Figure 12 shows sufficient reduction of fluctuations at 20 CSR simulations per cell. Finally, nearest neighbour distributions and complex formation metrics can be calculated analogously to the experimental data. Figure 5, Figure 6 and Supplementary Figures 6 and 10 show results from > 6 cells per cell line, thus also displaying cell to cell variability. To increase sensitivity in comparing measured results and simulated CSR simulations we display results of two representative cells per cell line (Supplementary Figures 7,8,9 and 11). By additionally plotting the 20 individual CSR simulations and the 20 bootstrapped samples of a given cell, precisions induced by finite sample size are directly visible. Thus, allowing for a reliable assessment whether the measured outcome is in agreement with a CSR scenario (within the given precision) or not.

### Figures

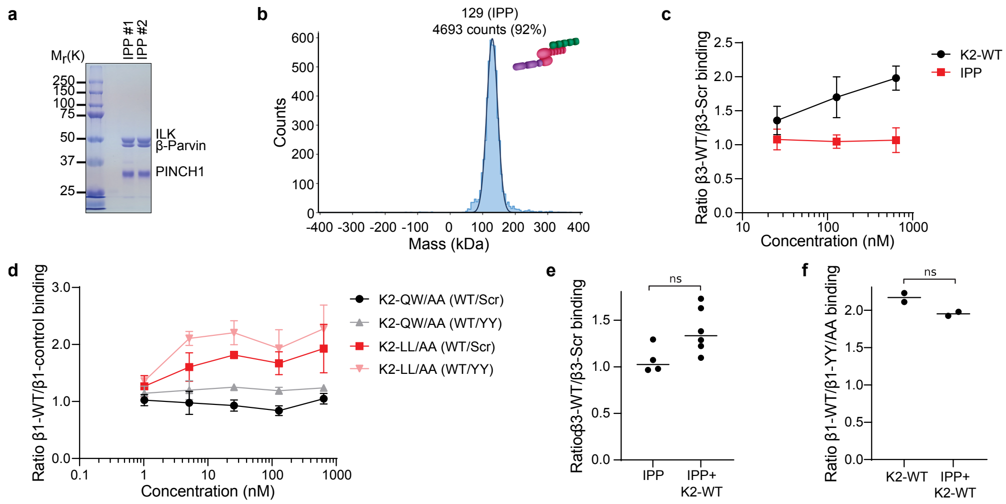


**Fig. S1. Purification of full-length IPP complex and interaction analyses with β integrin cytoplasmic tails. a** Purified IPP complex is resolved by SDS-PAGE and stained with Coomassie blue to show the high level of purity. **b** Mass photometry mass distribution of the purified, recombinant IPP complex. Histogram shows the particle counts of recombinant IPP complexes at the indicated molecular mass. **c** Quantification of the relative binding of K2-WT or IPP to β3-WT tail peptides compared to scrambled β3 tail peptides over the indicated concentration range (n=3 independent experiments; error bars indicate SD). **d** Quantification of the relative binding of K2-QW/AA and K2-LL/AA to β1-WT tail peptides compared to two control peptides, β1-YY/AA or scrambled β1-tail peptides, over the indicated concentration range (n=3 independent experiments; error bars indicate SD). **e** Increased binding of the IPP complex to β3 tail peptides after co-incubation with wild-type kindlin-2 (n=4-6 independent experiments, ns = not significant). **f** Presence of the IPP complex does not affect kindlin binding to β1-WT tail peptides. 25 nmol K2-WT was applied to immobilized β1-WT, β1-YY/AA, or scrambled β1-tail peptides either alone or in combination with 250 nM IPP complex. After removing unbound proteins, remaining proteins were detected with antibodies against kindlin-2 (n=2 independent experiments, ns = not significant).

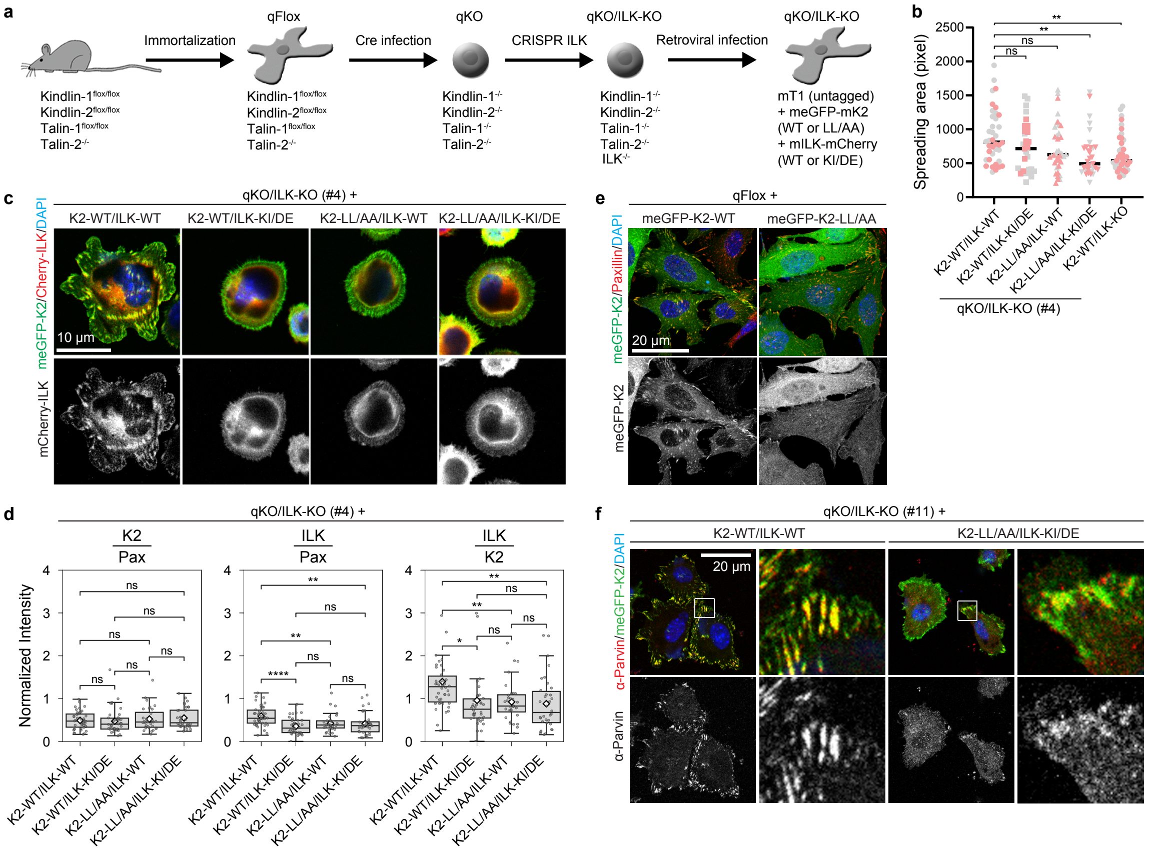


**Fig. S2. Cell system to study IPP recruitment to FAs. a** Scheme for the generation of talin/kindlin and ILK-deficient (qKO/ILK-KO) fibroblasts. ILK was depleted in talin and kindlin-deficient (qKO) fibroblasts by CRIPSR and two independent qKO/ILK-KO clones were subsequently retrovirally transduced with talin-1 and a combination of mEGFP-kindlin-2 (WT or LL/AA mutant) and mCherry-ILK (WT or KI/DE). **b** Quantification of the cell area of the indicated cell lines after spreading on FN for 30 min (>50 cells counted in three independent experiments; *t*-test significances are indicated; ** $0.001 < p \leq 0.01$). **c** Fluorescence images of mEGFP-K2 and mCherry-ILK in talin1- reconstituted qKO/ILK-KO fibroblasts spread for 30 min on FN. DAPI was used to stain nuclei. **d** FA-localized fluorescence intensities of mEGFP-K2 and mCherry-ILK measured in cells seeded for 75 min on FN. Paxillin servd as adhesion marker. Kindlin-2 intensities were normalized with respect to paxillin. ILK intensities were normalized relative to paxillin or to K2. Welch's *t*-test was applied to normalized intensities: ns (not significant) $p > 0.05$, * $0.01 < p \leq 0.05$, ** $0.001 < p \leq 0.01$, **** $p \leq 0.0001$. **e** Fluorescence images of qFlox cells expressing mEGFP-K2-WT or mEGFP-K2-LL/AA 75 min after plating on FN and stained with antibodies against paxillin (red). DAPI was used to stain nuclei. **f** Fluorescence images of mEGFP-K2 and mCherry-ILK in talin-1 expressing qKO/ILK-KO cells seeded for 75 min on FN and stained with antibodies against endogenous α-parvin. DAPI was used to stain nuclei.

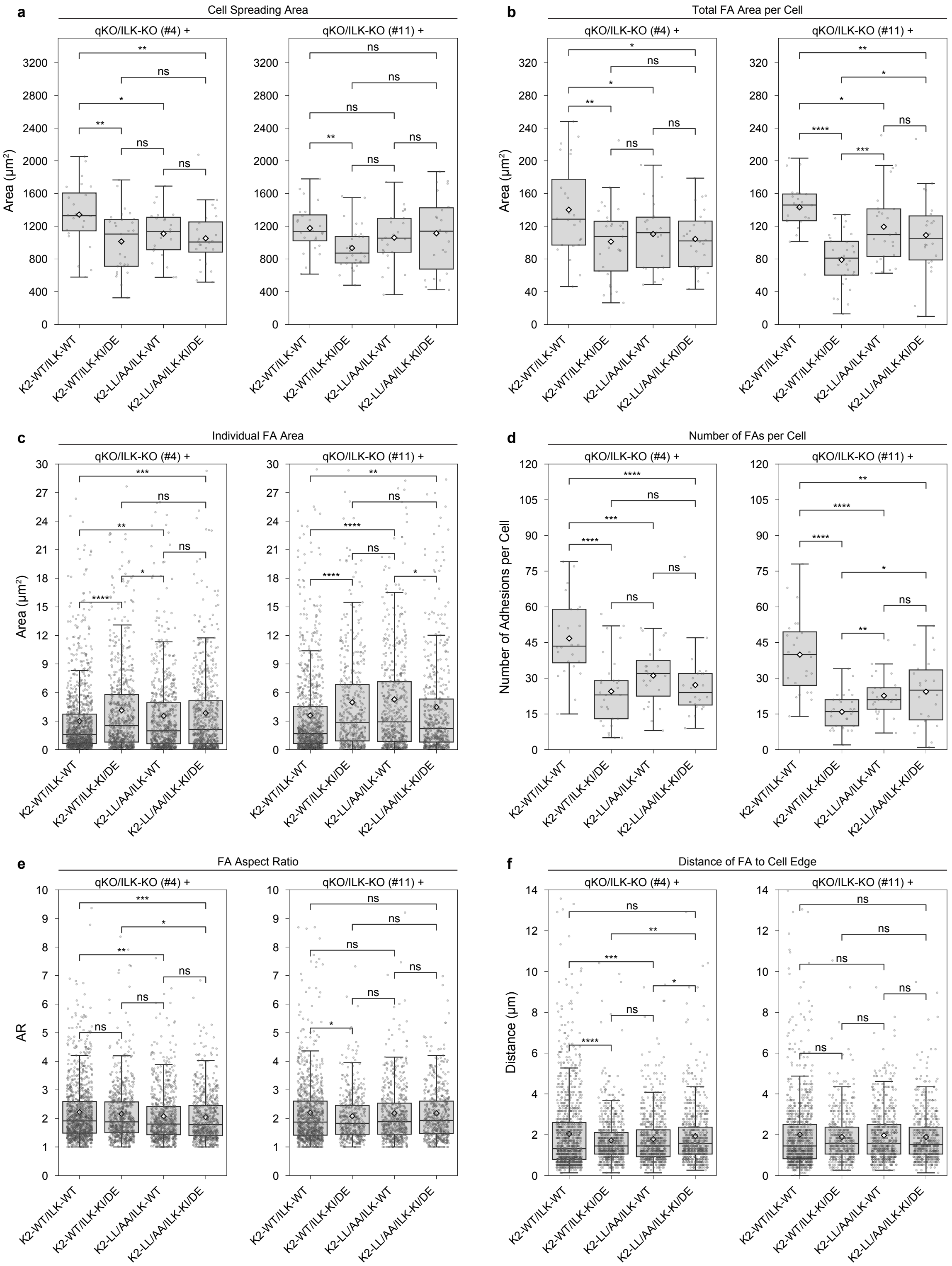


**Fig. S3. Focal Adhesion Morphology.** Quantification of FA morphology in cells seeded for 75 min on FN and derived from qKO/ILK-KO clone #4 and clone #11, respectively, reconstituted with talin1, mEGFP-K2 (WT or LL/AA) and mCherry-ILK (WT or KI/DE). Binary masks of the cell outline and FAs are based on K2 and paxillin signals, respectively, and were used to calculate morphology metrics. **a** Cell spreading area. **b** Total area of FAs per cell. **c** Area of individual FAs. **d** Number of FAs per cell. **e** Aspect ratio of individual FAs. **f** Distance of the FA centroids to the cell edge. For panels a, b, and d, $N \geq 22$ cells per condition were analysed, excluding cells touching the edge of the field of view. Data in panels c, e, and f represent $N \geq 650$ FAs per condition from $N \geq 64$ cells, including those touching the FOV edge. Boxplots show median and 0th and 75th percentile with whiskers reaching the last data point within 1.5 x interquartile range, white diamonds in the boxplots indicate mean. Welch's t-test at $\alpha = 0.05$ with ns (not significant) $p > 0.05$, * $0.01 < p \leq 0.05$, ** $0.001 < p \leq 0.01$, *** $0.0001 < p \leq 0.001$, **** $p \leq 0.0001$.

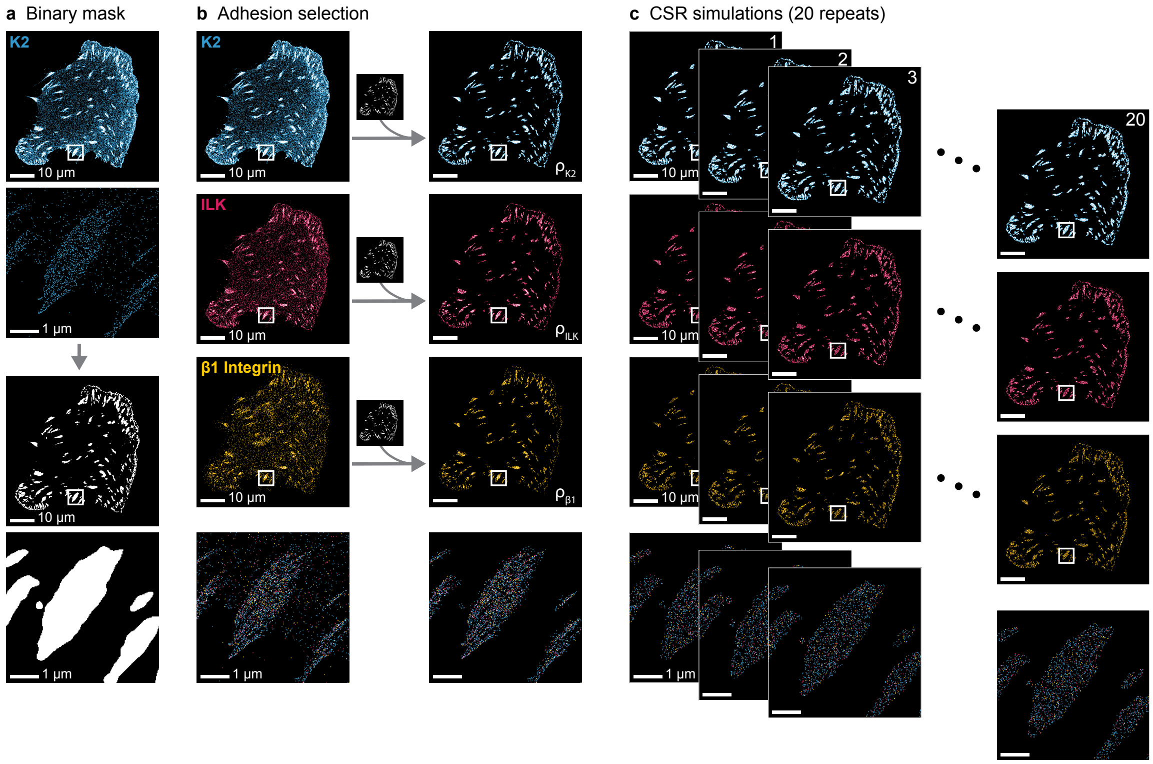


**Fig. S4. Identification of FAs and CSR simulations**. **a** FAs are identified using a binary mask (bottom, white: FA, black: non-FA area) based on the K2 protein coordinates (top). **b** K2, ILK and active β1 integrin proteins within FAs are selected using the generated mask. **c** CSR simulations are performed for each target within the masked area based on measured protein densities. This allows to compare in situ protein patterns to a simulated scenario of non-interacting proteins. Densities are based on the number of proteins within the FA area identified by the mask. Fluctuations due to finite protein numbers are reduced in CSR simulations by repeating the simulation 20 times, calculating protein interaction metrics for each repeat and taking the average over all CSR repeats of a given cell.

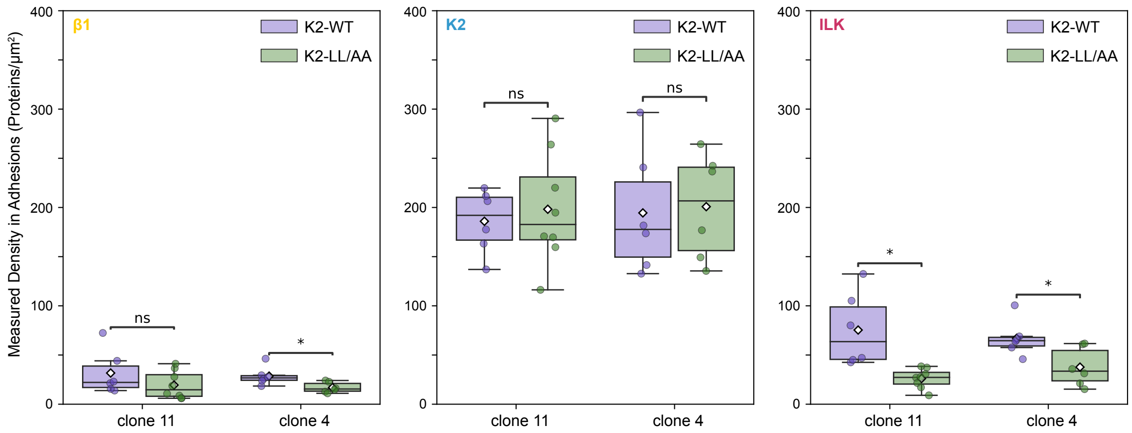


**Fig. S5. Density of ILK, K2 and active β1 integrins in FAs**. The FA surface area for protein density measurements is determined via a binary mask (see Methods and Supplementary Figure 4). Labelled single molecules within the FA mask are identified to calculate the apparent surface density of active β1 integrins, K2 and ILK. Active β1-integrin densities are slightly reduced in K2-LL/AA expressing cells compared to the respective K2-WT cells in clone 4, but not in clone 11. The density of K2-LL/AA is unaffected in FAs. The density of ILK is significantly reduced in FAs when co-expressed with K2-LL/AA. Boxplots show median and 0th and 75th percentile with whiskers reaching the last data point within 1.5 x interquartile range, white diamonds in the boxplots indicate mean. Datapoints represent protein densities in FAs of individual cells (N ≥ 6 cells per cell line). Welch's t-test at $\alpha = 0.05$ with ns (not significant) $p > 0.05$, * $0.01 < p \leq 0.05$.

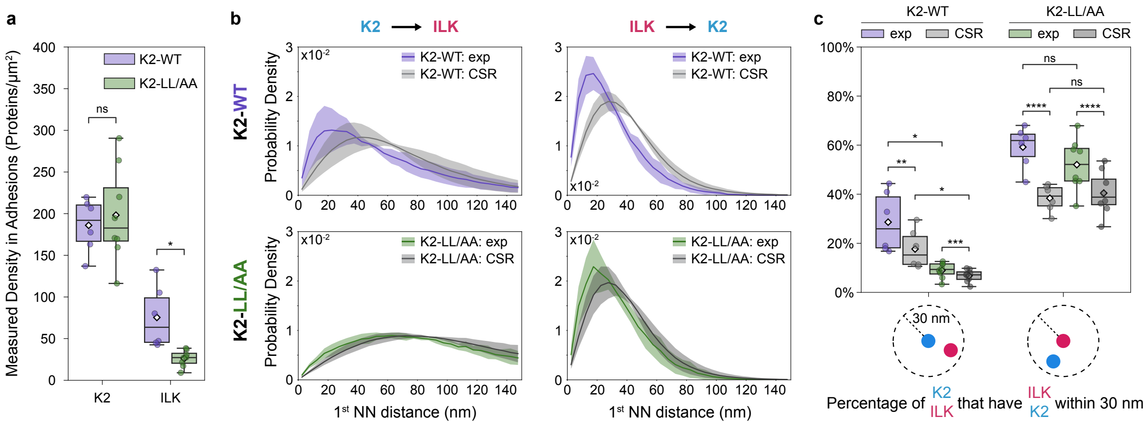


**Fig. S6. K2-LL/AA mutation decreases the proximity to ILK in FAs (clone 11)**. **a** Observed surface density of K2 and ILK molecules in FAs. The density of K2 remains unaffected K2-LL/AA expressing cells, while ILK density is significantly reduced in K2-LL/AA cells compared to K2-WT. Reported values represent observed densities and do not account for labelling efficiency. Welch's t-test at $\alpha = 0.05$; ns (not significant) $p > 0.05$, * $0.01 < p \leq 0.05$. **b** Average nearest neighbour distributions between K2 and ILK molecules in FAs are compared to a scenario simulating complete spatial randomness (CSR). Left (K2 → ILK): ILK is significantly closer to K2 than in the CSR simulation representing the absence of molecular interaction ($p < 0.0001$). The K2-LL/AA mutation results in an average NND distribution closer to the CSR scenario, but remains statistically different from the CSR scenario ($p < 0.0001$). Measured K2-WT and LL/AA curves differ ($p < 0.0001$). Right (ILK → K2): Experimental distributions differ significantly from CSR in both K2-WT ($p < 0.0001$) and K2-LL/AA cells ($p < 0.0001$), while the K2-LL/AA is more similar to CSR. Modified Chi-squared test according to Ref.(9) at $\alpha = 0.05$. **c** Percentage of K2 molecules with at least one ILK molecule within 30 nm (left) and vice versa (right) in FAs compared to CSR. The percentage of K2 with nearby ILK drops from WT to mutant cells and closely approaches CSR in mutant cells, with the deviation from CSR remaining significant. ILK with nearby K2 does not show the significant drop from K2-WT to K2-LL/AA cells and remains significantly elevated over its CSR baseline in K2-LL/AA expressing cells. Boxplots show median and 25th and 75th percentile with whiskers reaching the last data point within 1.5 x interquartile range, diamonds indicate mean. Paired two-sample t-test for exp vs. CSR and Welch's t-test for K2-WT vs. K2-LL/AA. ns (not significant) $p > 0.05$, * $0.01 < p \leq 0.05$, ** $0.001 < p \leq 0.01$, *** $0.0001 < p \leq 0.001$, **** $p \leq 0.0001$. Quantifications based on N = 6 cells for mCherry-ILK-WT/mEGFP-K2-WT and N = 8 for mCherry-ILK-WT/mEGFP-K2-LL/AA

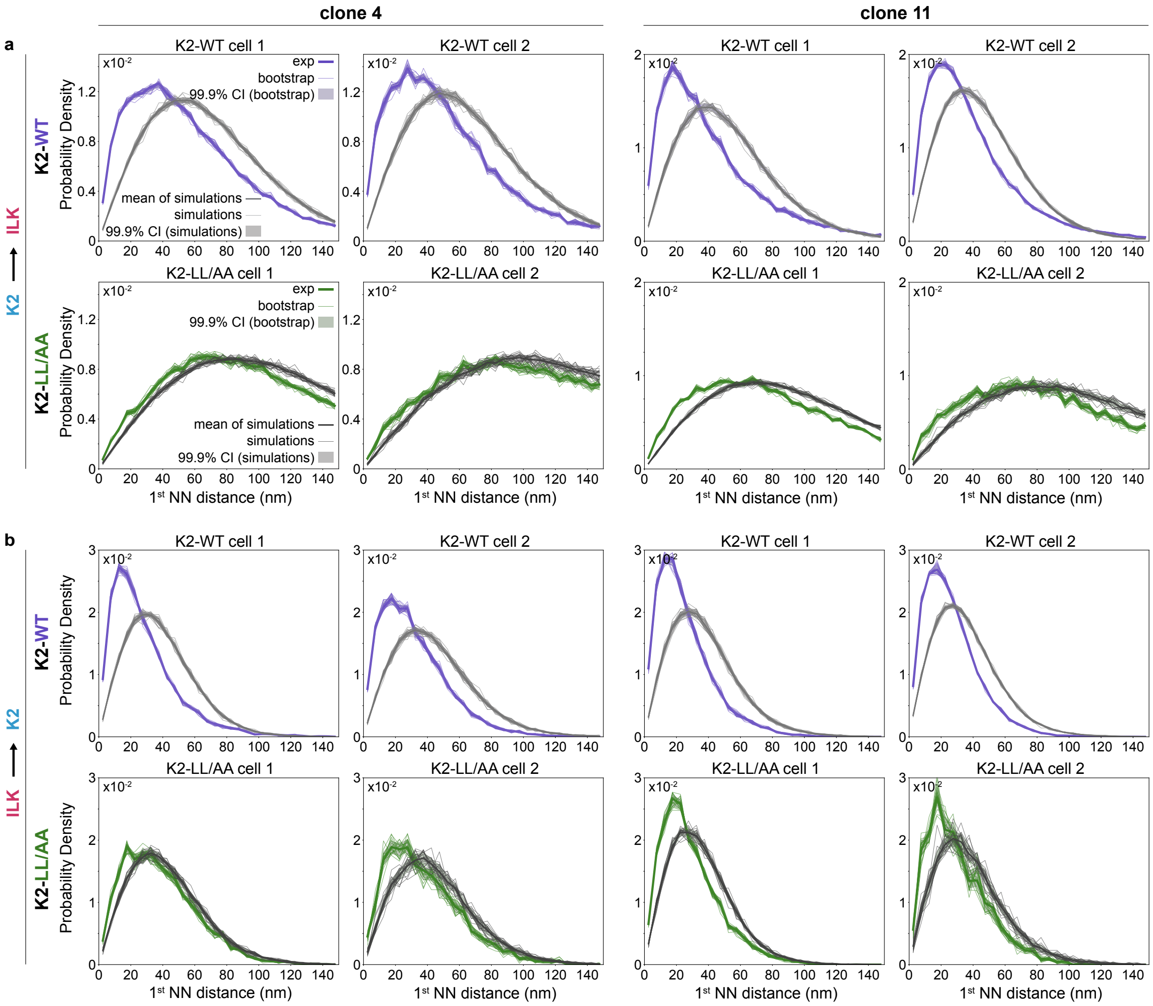


**Fig. S7. Single cell nearest neighbour distance distributions between K2 and ILK.** Nearest neighbour distance distributions in FAs of two representative cells per clone 4 and 11, and cells expressing mCherry-ILK-WT together with either mEGFP-K2-WT or mEGFP-K2-LL/AA. Each plot shows the measured NND distribution ("exp") and the simulated CSR NND distribution ("mean of simulations", based on 20 simulations) for a given cell. Additionally, 20 bootstrapped samples of the measured NND and 20 CSR simulations are shown to visualize the precision given by the finite number of molecules per cell. **a** The NND distributions from K2-WT to ILK (top) differ from the respective CSR simulations. The NND distributions of K2-LL/AA to ILK (bottom) approach the CSR scenario suggesting that the mutation almost fully prevents binding to ILK. However, short distances NNDs below around 60 nm are enlarged compared to the CSR simulation. Notably, the bootstrapped samples of the measured NNDs and the individual CSR distributions do not overlap in this region, indicating that there are remaining molecular interactions. **b** The NND distributions from ILK to K2-WT and K2-LL/AA for the same cells as in (a) show a difference to the CSR scenario. Notably, the NNDs from ILK to K2 show a more pronounced deviation from CSR than the NNDs from K2 to ILK in panel (a). Kolmogorov-Smirnov test between the measured NNDs and NNDs from individual CSR simulations as well as the mean of all simulations always yields $p < 0.0001$.

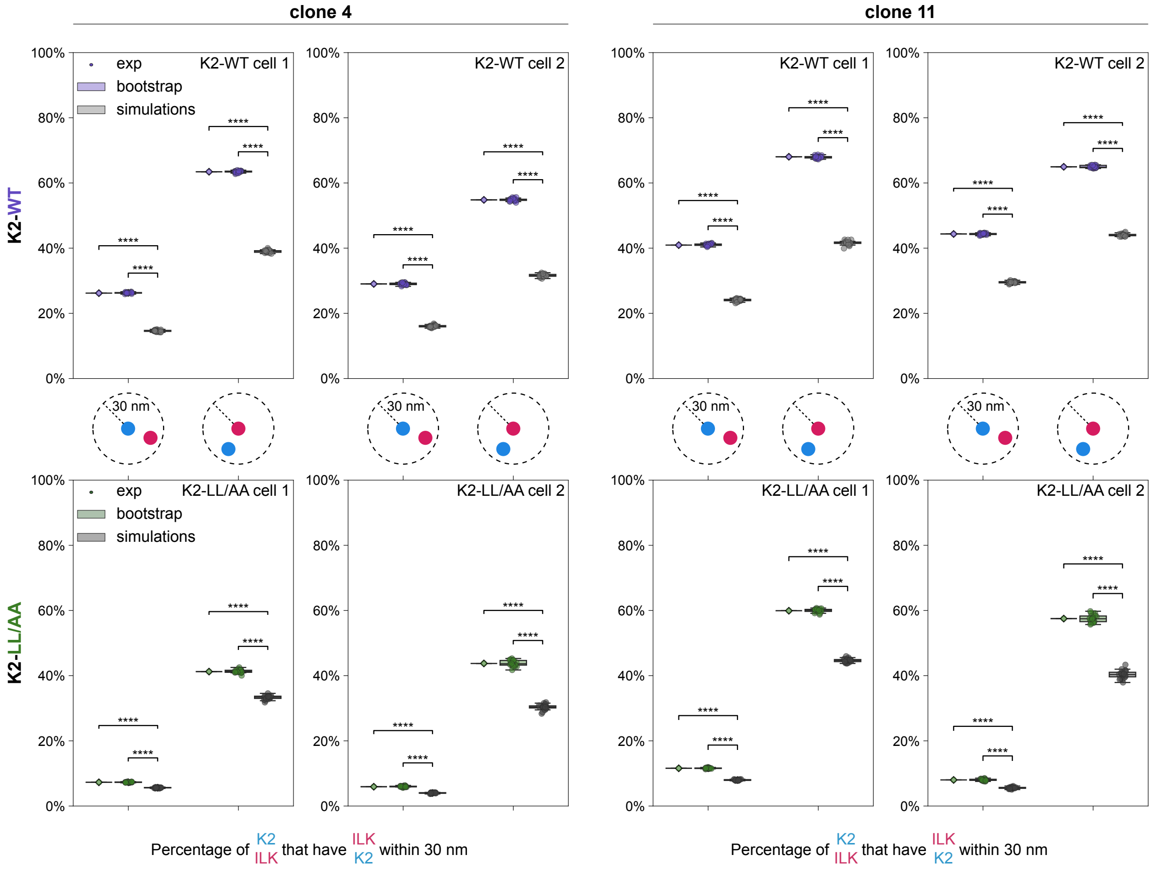


**Fig. S8. Single cell colocalization of K2 and ILK.** Percentage of K2 molecule with at least one ILK molecule within 30 nm and vice versa in FAs compared to CSR for two representative cells per clone (clones 4 and 11) and cell line (expressing mCherry-ILK with mEGFP-K2-WT or mEGFP-K2-LL/AA). Shown are the experimentally measured percentage, values from 20 bootstrap samples and 20 CSR simulations respectively. Boxplots show median and 25th and 75th percentile with whiskers reaching the last data point within 1.5 x interquartile range, diamonds indicate mean. Two-sample t-test for bootstrap samples vs. CSR simulations and one-sample t-test for exp vs. CSR simulations: **** $p \leq 0.0001$.

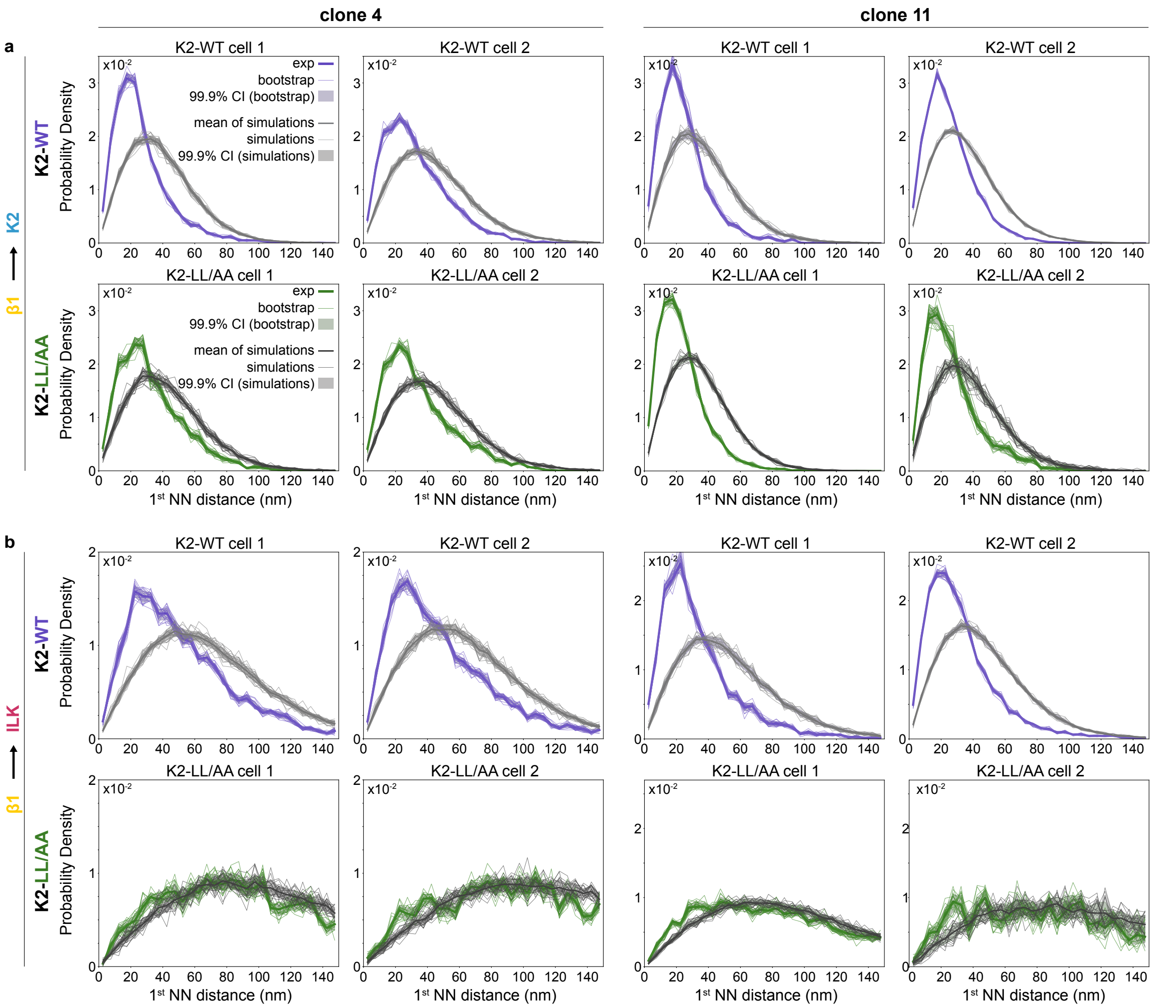


**Fig. S9. Single cell nearest neighbour distance distributions between β1 and K2 and β1 and ILK.** Nearest neighbour distance distributions in FAs of two representative cells per clone 4 and 11, and cells expressing mCherry-ILK-WT together with either mEGFP-K2-WT or mEGFP-K2-LL/AA. Each plot shows the measured NND distribution ("exp") and the simulated CSR NND distribution ("mean of simulations", based on 20 simulations) for a given cell. Additionally, 20 bootstrapped samples of the measured NND and 20 CSR simulations are shown to visualize the precision given by the finite number of molecules per cell. **a** The NND distributions from β1 to K2-WT and K2-LL/AA for the same cells as in **Fig. S7** and **S8**, show a clear difference to the CSR scenario. **b** The NND distributions from β1 to ILK-WT for the same cells as in (a). In K2-WT expressing cells the measured distribution and the CSR simulation differ indicating recruitment of ILK-WT to β1. In K2-LL/AA cells the distance of ILK to β1 is comparable to the CSR results, with only a small amount of ILK proteins closer to β1 than expected from a CSR scenario. Kolmogorov-Smirnov test between the measured NNDs and NNDs from individual CSR simulations as well as the mean of all simulations always yields $p < 0.0001$.

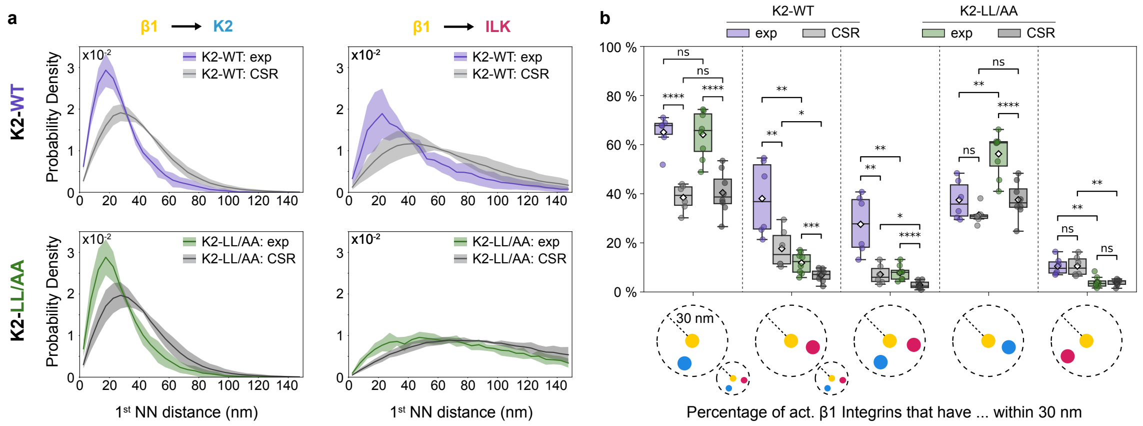


**Fig. S10. ILK recruitment to β1 integrins is fully dependent on K2 (clone 11). a** Nearest neighbour distances in FAs between active β1-integrin and K2 or ILK are compared to simulated CSR scenarios (Mean ± SD of N NND distributions from N cells per cell line). Left (β1 → K2): In K2-WT cells, K2 is significantly closer to active β1-integrins (top left) than in absence of molecular interaction ($p < 0.0001$). This equivalently holds true for K2-LL/AA ($p < 0.0001$). K2-LL/AA and K2-WT distributions do not differ ($p = 0.99$ for measured experimental NND distributions and $p = 0.99$ for simulated CSR distributions), demonstrating that the LL/AA mutation does not significantly change K2's interaction with active β1-integrins. Right (β1 → ILK): The experimental and CSR distributions in K2-LL/AA cells are shifted to larger distances compared to K2-WT ($p < 0.0001$ respectively). ILK is significantly closer to active β1-integrins in K2-WT expressing cells ($p < 0.0001$). In K2-LL/AA expressing cells, the measured curve is more similar to the CSR scenario, but still significantly different ($p < 0.0001$). Modified Chi-squared test according to Ref.(9) at $\alpha = 0.05$. **b** The percentage of active β1 integrins that have a K2 molecule, an ILK molecule, K2 and ILK simultaneously or only one of them within a radius of 30 nm around them are calculated. Boxplots show median and 25th and 75th percentile with whiskers reaching the last data point within 1.5 x interquartile range, diamonds indicate mean. Paired two-sample t-test for exp vs. CSR and Welch's t-test for K2-WT vs. K2-LL/AA: ns (not significant) $p > 0.05$, * $0.01 < p \leq 0.05$, ** $0.001 < p \leq 0.01$, *** $0.0001 < p \leq 0.001$, **** $p \leq 0.0001$. Quantification based on N = 6 cells for mCherry-ILK-WT/mEGFP-K2-WT and N = 8 for mCherry-ILK-WT/mEGFP-K2-LL/AA

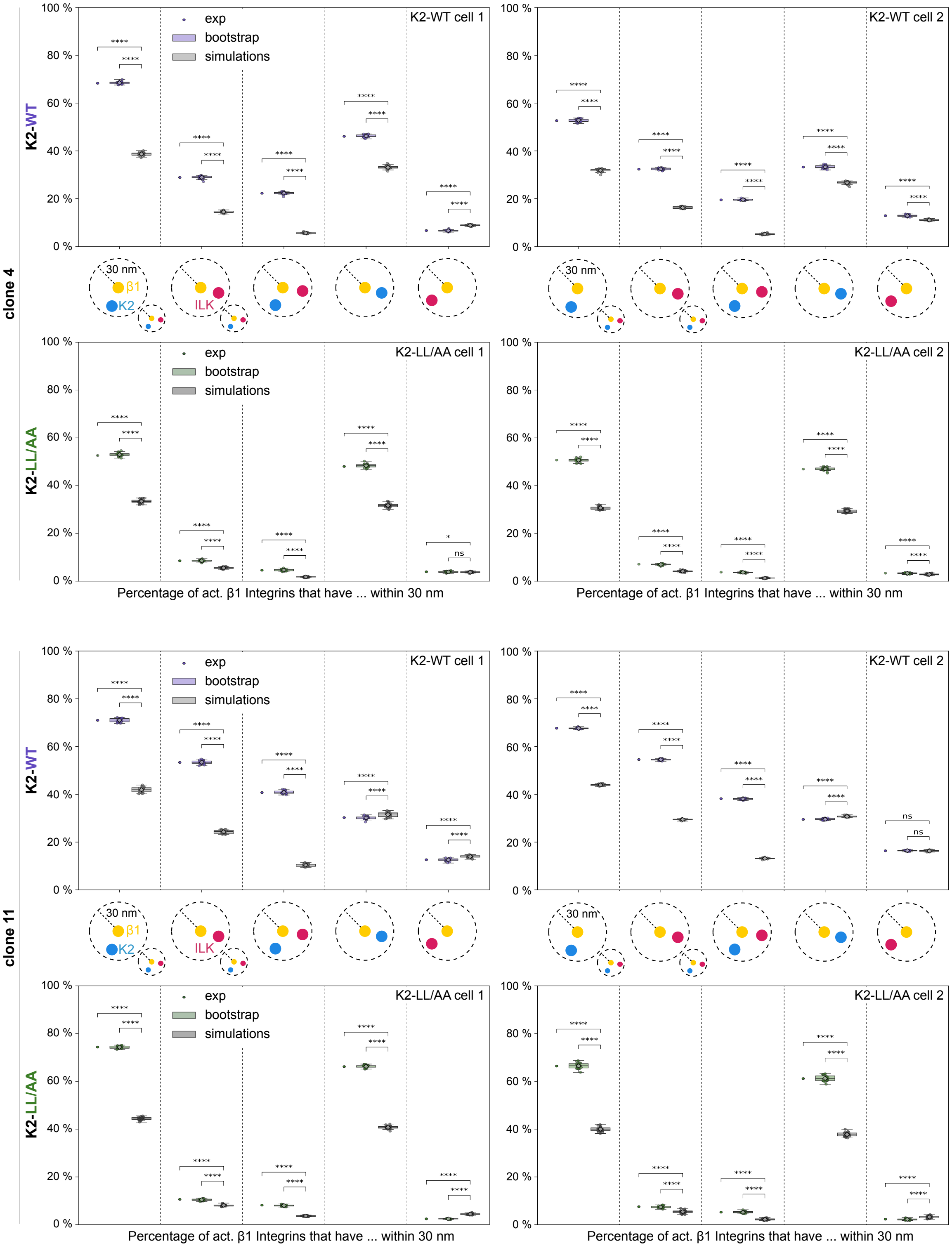


**Fig. S11. Single cell colocalization of K2 and ILK near active β1 integrins.** Percentage of active β1 integrins that exclusively or simultaneously colocalize with K2 and ILK within a radius of 30 nm for two representative cells per clone (clones 4 and 11) and cell line (expressing mCherry-ILK with mEGFP-K2-WT or mEGFP-K2-LL/AA). Shown are the experimentally measured percentage, values from 20 bootstrap samples and 20 CSR simulations respectively. Boxplots show median and 25th and 75th percentile with whiskers reaching the last data point within 1.5 x interquartile range, diamonds indicate mean. Two-sample t-test for bootstrap samples vs. CSR simulations and one-sample t-test for exp vs. CSR simulations: ns (not significant) $p > 0.05$, **** $p \leq 0.0001$.

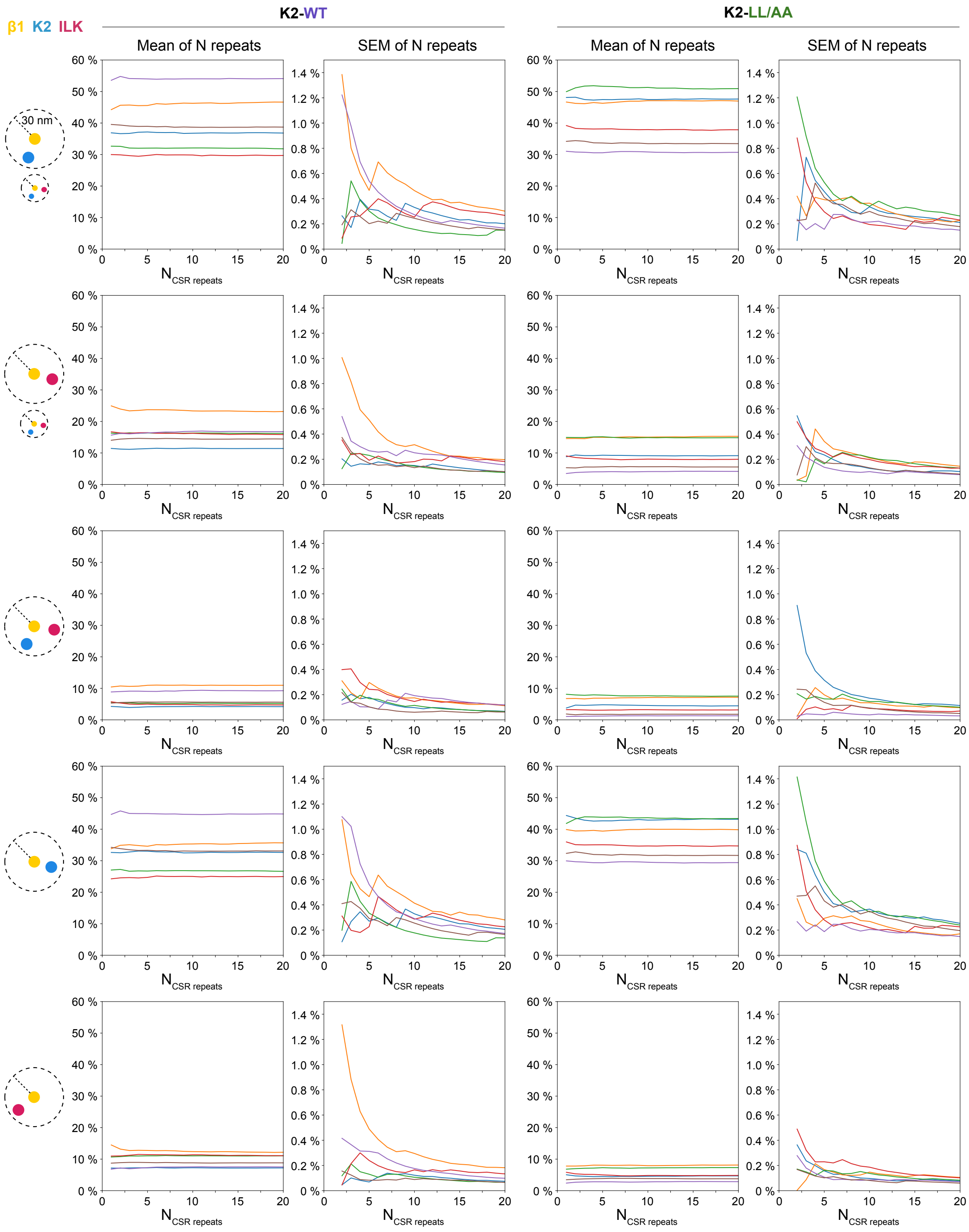


**Fig. S12. Increasing precision in CSR simulations.** For each cell the CSR simulation is performed with the experimentally determined number of molecules, thus limited in its precision by the finite sample size. By repeating the simulation and proceeding with the mean of multiple simulations the precision of any output metric is increased. Here we plot how the percentages of β1 integrins that colocalize with either K2, ILK, K2 & ILK, or only K2 or only ILK converge, when taking the mean over an increasing number of CSR simulations. Additionally, we display the standard error of the mean. Each curve corresponds to one cell of clone 4.
